# Core Bacterial and Host Fruit-Specific Yeast Microbiota in a Polyphagous Fly Pest

**DOI:** 10.64898/2026.04.24.720762

**Authors:** Sido Dunis, Marie Lapègue, Candice Deschamps, Lily Cesari, Anne Loiseau, Benoit Facon, Nicolas O. Rode

**Author notes:** Address correspondence to Sido Dunis, and Nicolas O. Rode.

## Abstract

Holometabolous polyphagous insects undergo metamorphosis and exploit multiple host plants, exposing them to highly variable ecological conditions across both life stages and host plants. Whether these insects harbour a stable core microbiota, a life stage- or a host plant-specific microbiota remain open questions. Here, we characterised fungal and bacterial communities associated with *Drosophila suzukii* across life stages (larvae, pupae and emerging flies) and host fruits (cherry, blackberry and strawberry) using 16S and ITS metabarcoding. We tested the relative influence of life stage and host fruit on microbiota composition using community and network-based analyses. First, we identified that yeasts, but not bacteria, were strongly structured among host fruits. Bacteria and filamentous fungi were shared across fruits, constituting a fruit core microbiota dominated by *Gluconobacter cerinus, Tatumella* sp. and *Cladosporium* spp. In contrast, yeast were rather fruit-specific: *Hanseniaspora* and *Pichia* genera mostly associated with cherries and strawberries, contrary to *Metschnikowia* with blackberries. Second, we found that fungal, but not bacterial, communities of *D. suzukii* were structured among host fruits. *D. suzukii* harboured a core bacterial microbiota composed of *G. cerinus* and a niche-specific (host plant-specific) microbiota composed of yeasts: *Hanseniaspora* typical in individuals related to cherry and strawberry, and *Metschnikowia* to blackberry. We also verified that individuals from artificial laboratory infestations and natural field infestations harboured similar microbial communities when they developed under the same laboratory conditions. Components of both core and niche-specific microbiota were most likely horizontally acquired by *D. suzukii* from host fruits. Taken together our results underline the importance of metacommunity approaches to investigate tripartite interactions among insects, host plants and microbiota.

**IMPORTANCE:** The role of microbiota in mediating interactions between phytophagous insects and their host plants has been well illustrated in specialist species. However, it has been less comprehensively studied in polyphagous species, which infest multiple host plants, and across life stages for holometabolous species experiencing separate ecological niches through development. We tested the existence of a core, a niche-specific and a stage-specific microbiota in a polyphagous holometabolous species, *D. suzukii*. We examined both fungal and bacterial communities in larvae, pupae and emerging flies infesting three host fruits. Our results showed first that the assembly of bacteria, filamentous fungi and yeasts on fruits is driven by different ecological processes. Second, that *D. suzukii* harbours a core bacterial microbiota, a niche-specific microbiota constituted by yeasts and no stage-specific microbiota. Our study emphasizes the importance of considering jointly the assembly of host plant and polyphagous insect microbial communities to better understand insect-microbe interactions.

## INTRODUCTION

Microbial symbionts play a central role in mediating interactions between phytophagous insects and their host plants (1,2). By influencing nutrient acquisition, detoxification, immunity and development, they can substantially affect insect fitness (3–8). Insects harbour very diverse microbial communities in structure and function in their digestive system (9,10). While insect-bacteria symbiosis has been extensively studied, fungal communities are increasingly recognized as important components of the microbiota but remain rarely studied jointly with bacteria (11–13).

Microbiota assembly is shaped by transmission (vertical versus horizontal), ecological filtering by the host or environment, and developmental remodeling (14–16). Within these microbial communities, the core microbiota, defined as taxa consistently associated with a given host genotype across time and space, is hypothesized to gather the most ecologically and functionally important species (17). Specialist phytophagous insects, associated with a single niche, usually harbour a core microbiota (18). For instance, *Bactrocera oleae* is strictly and constantly associated with *Candidatus* Erwinia dacicola (19). In contrast, polyphagous insects experience heterogeneous nutritional environments across host plants, potentially favoring flexible and environmentally acquired microbiota. Metamorphosis adds a further layer of complexity, as gut remodeling affects microbial abundance and diversity (20–24), life stages may harbour distinct communities due to differing ecological niches and nutritional requirements (25). Thus, polyphagous holometabolous insects face the challenge of maintaining beneficial microbial functions across both host plant variability and developmental transitions.

Three non-exclusive scenarios can be envisioned: (i) a persistent core microbiota shared across life stages and hosts; (ii) a stage-specific microbiota driven by developmental processes; and (iii) a niche-specific microbiota structured by host plants (Fig. 1). These scenarios imply different modes of microbial acquisition (vertical and/or horizontal transmission) and varying degrees of association stability (transient or persistent). The relative importance of these scenarios remains poorly characterized in polyphagous holometabolous insects.

**Figure 1:**
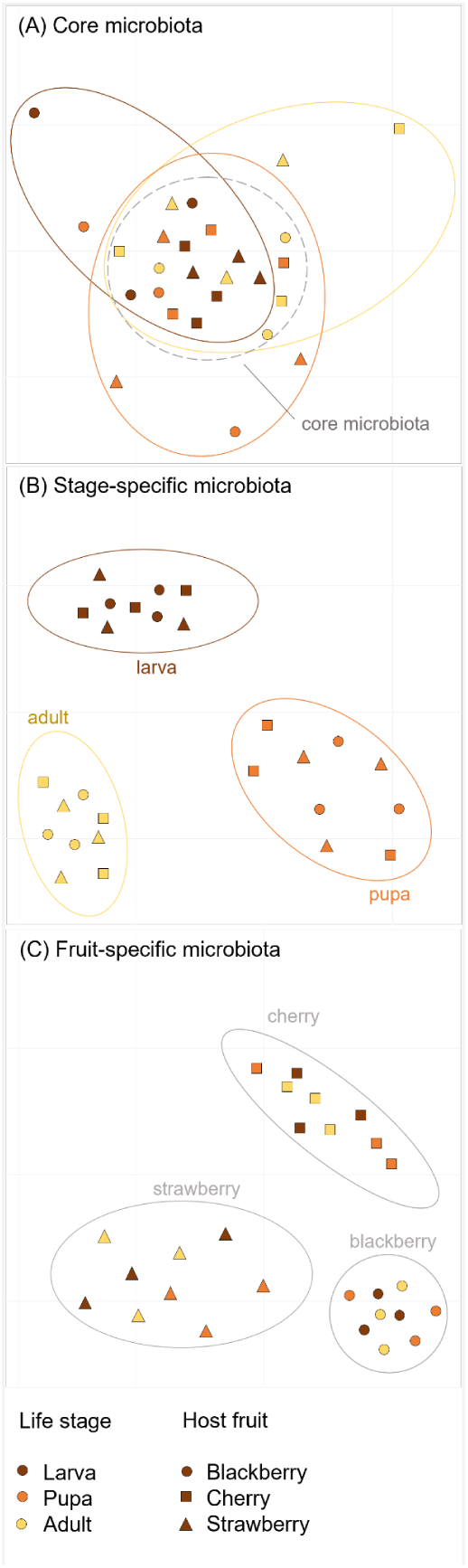
Predicted ordination plot of *D. suzukii* microbiota under the hypothesis of (A) a core microbiota shared across life stages and host fruits, (B) a stage-specific microbiota structured among life stages and (C) a fruit-specific microbiota structured among host fruits.

*Drosophila* is a well-established model for microbiota studies. Although microbial variations across life stages have been documented under laboratory conditions (22,26,27), they remain poorly characterized in natural contexts and across different host plants. Shifts in microbial communities across distinct habitats have been documented in adults (28–30), but such variation has not been examined through the full life cycle or jointly across bacterial and fungal communities. In particular, there is a lack of data regarding differences in larval and pupal microbiota as the few studies considering larvae have not compared their microbiota across hosts (31–33). By examining microbial communities across host plants and Drosophila life stages, metacommunity-level analyses can help identify the ecological processes underlying observed community patterns – including selection imposed by host plants or developmental stages, dispersal, stochastic drift, and diversification (34).

Here, we investigated fungal and bacterial communities of *Drosophila suzukii* across life stages and host fruits. This invasive pest with a broad host plant range (>80, (34)) is particularly relevant given larval dependence on fruit-associated microbes (4). We sampled blackberries, cherries, and strawberries from fields, allowed oviposition by lab-reared flies, and characterized bacterial and fungal communities using 16S and ITS metabarcoding. We tested three non-mutually exclusive hypotheses: (i) the presence of a core microbiota shared across life stages and host fruits; (ii) the existence of stage-specific microbial communities; and (iii) the occurrence of niche-specific communities structured by host fruit identity. Using community and network-based analyses, we evaluated the relative influence of life stage and host fruit on microbiota composition.

## RESULTS

### Variation in microbial communities composition over life cycle and host fruit

Larvae which developed in artificially or naturally infested fruits harboured the same major microbial families. Likewise, pupae and emerging adults which developed on fruit medium from both lab and wild larvae harboured the same major microbial families (see Fig. S5 in supplemental material).

The number of bacterial and fungal families harboured by *D. suzukii* decreased over the life cycle (Fig. 2). Acetobacteraceae and Lactobacillaceae were the two main bacterial families in lab flies (Fig. 2A, “Mothers”). After 48 hours on fruits, mothers also harboured Erwiniaceae, the most prevalent family in uninfested fruits (Fig. 2A, “Fruits”). These families remained dominant in larvae and host fruits after 5-6 days, regardless of fruit type. Acetobacteraceae appeared on fruits after 5-6 days, being infested or not (see Fig. S6A in supplemental material), and became dominant in later life stages (pupae, emerging adults). The diversity of bacterial genera within major families was minimal (max. 2, see Fig. S7A in supplemental material).

**Figure 2:**
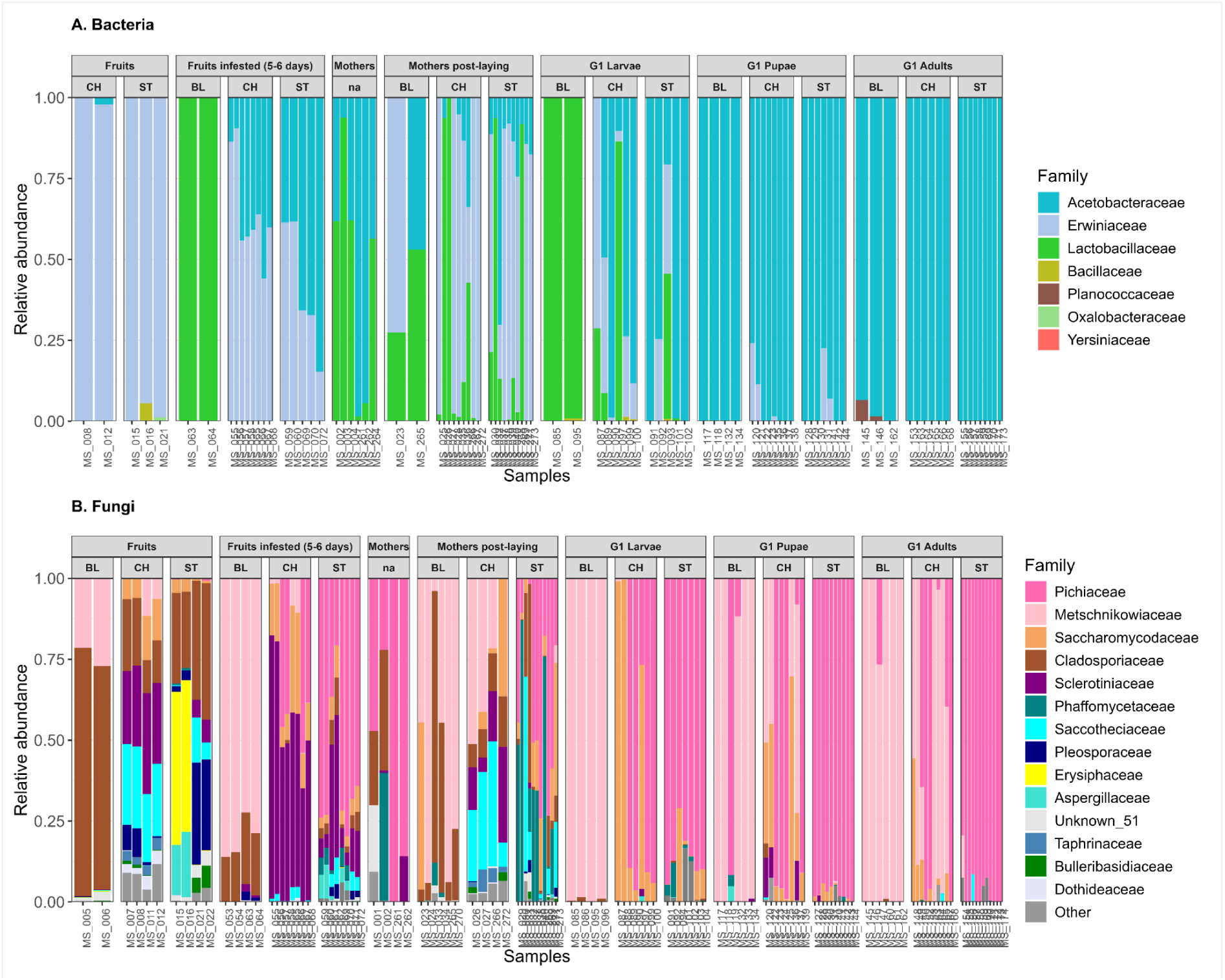
Relative abundance of the seven most abundant bacterial (A) and 15 fungal families (B) of *D. suzukii* individuals and their host fruits, for each combination of type of sample and host fruit. We sampled undamaged (”Fruits”) and artificially infested (“Fruits infested, 5-6 days”) host fruits, females lab flies used to infest the fruits before (“Mothers”) and after egg-laying on host fruits (“Mothers post-laying”), and the offspring at larval (“G1 Larvae”), pupal (“G1 Pupae”) and emerged adult (“G1 Adults”) stages. BL: blackberry, CH: cherry, ST: strawberry. Unknown_51 fungal ASV belongs to the Fungi kingdom based on BLAST, but its phylum is undetermined.

Fungal communities in lab flies were dominated by Pichiaceae, Cladosporiaceae and Phaffomycetaceae (Fig. 2B, “Mothers”). After fruit exposure, mothers also harboured additional fruit-associated taxa (e.g. Sclerotiniaceae, Pleosporaceae, Taphrinaceae; Fig. 2B, “Fruits”). After 5-6 days host fruits were dominated by Pichiaceae, Metschnikowiaceae or Saccharomycodaceae along with either Sclerotiniaceae or Cladosporiaceae (Fig. 2B, “Fruits infested (5-6 days)”). After 5-6 days, these dominant fungal families were found on both infested and uninfested fruits (see Fig. S6B in supplemental material). All *D. suzukii* life stages harboured at least one ASV of a single genus within the Metschnikowiaceae, Pichiaceae or Saccharomycodaceae families (respectively the *Metschnikowia, Pichia* or *Hanseniaspora* genera, see Fig. S7B in supplemental material).

### Diversity

Consistent with composition data, bacterial richness (observed richness and Chao1) and diversity (Shannon index) varied among the three types of fruits, and decreased between larvae, pupae and adults (Fig. 3A). Fungal richness and diversity also varied among fruits and decreased across life stages for strawberry, but not for cherry and blackberry (Fig. 3B). Samples from blackberry had both higher fungal richness and diversity than strawberry and cherry samples, while their bacterial richness and diversity was similar across life stages.

**Figure 3:**
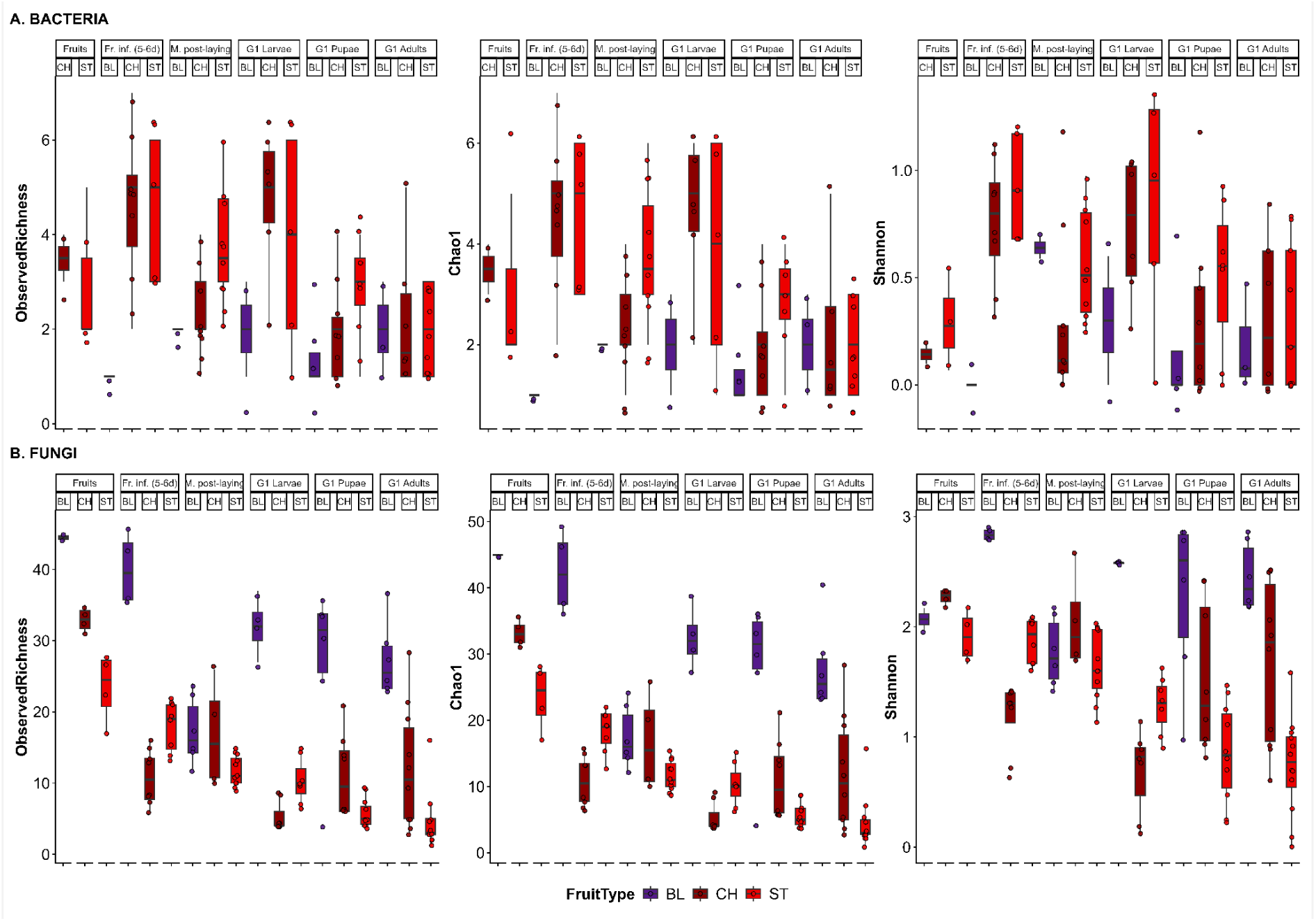
Alpha diversity (richness) indices for bacteria (A) and fungi (B) in *D. suzukii* individuals and their host fruits, grouped by sample type (undamaged (“Fruits”) and artificially infested host fruits (“Fruits infested, 5-6 days”), females lab flies used to infest host fruits after egg-laying (“Mothers post-laying”), and the offspring at larval (“G1 Larvae”), pupal (“G1 Pupae”) and emerged adult (“G1 Adults”) stages), and host fruit type (BL: blackberry, CH: cherry, ST: strawberry). For bacteria, blackberry samples had either fewer than 3,000 reads, a single or max. two bacterial ASVs. Mothers before egg-laying samples (“Mothers”) were removed from the analyses.

The richness and diversity of both bacterial and fungal communities of host fruits were similar between infested and uninfested fruits after 5-6 days (see Fig. S8B in supplemental material).

### NMDS

Nonmetric multidimensional scaling (NMDS) suggested that bacterial communities harboured by *D. suzukii* individuals showed some structure among sample types, rather than among host fruits (Fig. 4A). Samples clustered by developmental stages (colors), with Acetobacteriaceae presence correlated with both NMDS axes. Conversely, fungal communities were structured by host fruits (Fig. 4B). Samples clustered by host fruit (shapes), separating strawberry and cherry from blackberry on the NMDS1 axis. In line with these results and regardless of infestation status, the fungal communities of fruits were strongly structured among fruits and substructured among fruit varieties and sampling sites contrary to bacterial communities which were only slightly structured among varieties and sampling sites (Dataset 2, see Fig. S9 in supplemental material).

**Figure 4:**
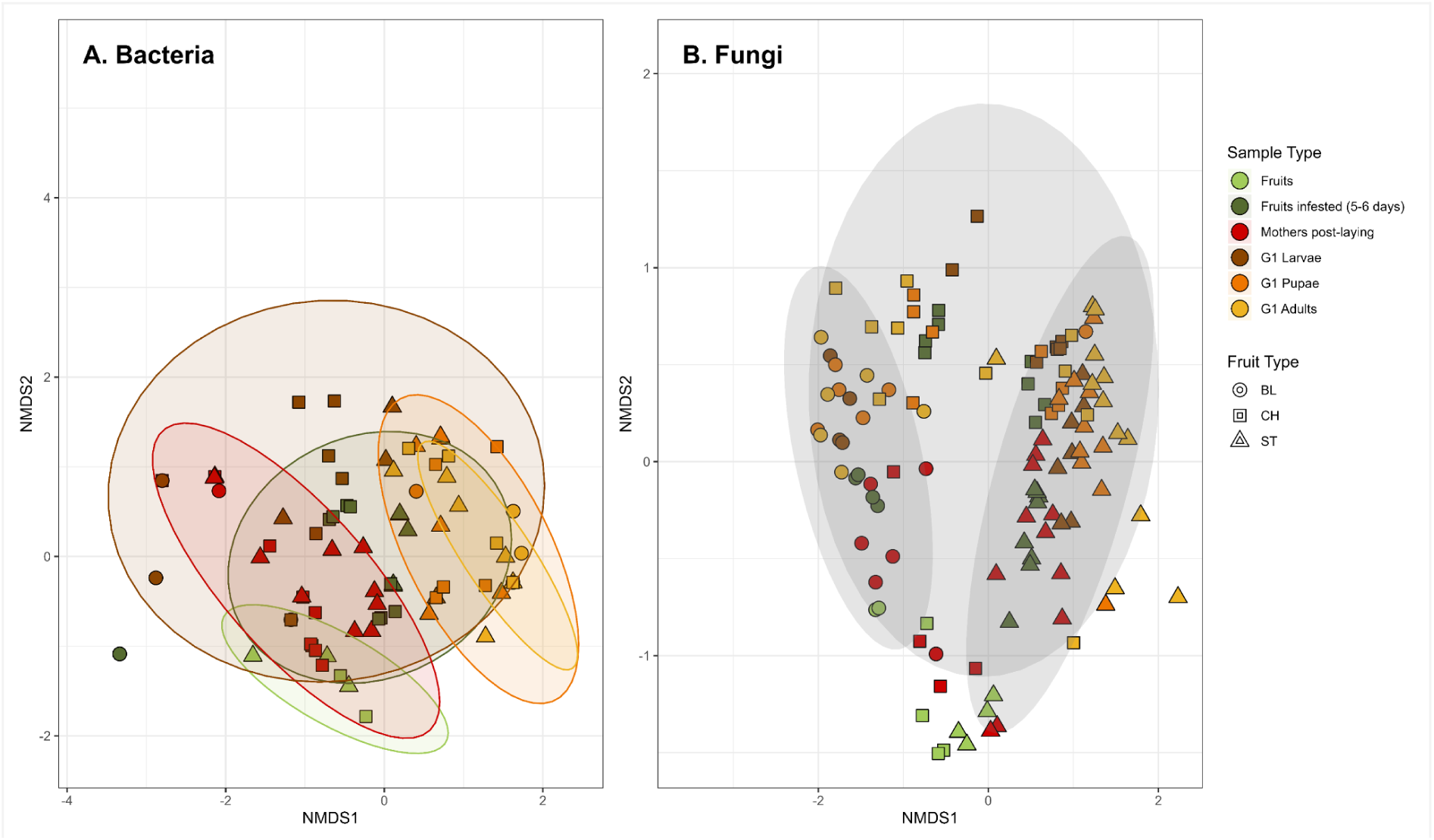
Non-metric multidimensional scaling (NMDS) ordination plots of bacterial (A) and fungal (B) communities of *D. suzukii* individuals and their direct host fruits based on Jaccard distance metric (stress = 0.1042 and 0.1616 respectively). A point represents a pool of samples (fruits or individuals). Symbols represent fruit type: BL = blackberry, CH = cherry, ST = strawberry. Colors represent sample types (mothers before egg-laying samples (Mothers), were pulled out as not related to host fruits). Ellipses are plotted by Sample Type for bacteria (A), and by Fruit Type for fungi (B). Sample ordinations based on Bray-Curtis distances were not different, indicating that microbial species’ abundances were balanced (see Fig. S10 in supplemental material).

### Network analysis

#### Sample groups

Latent Block Model (LBM) based on bacterial communities classified all samples in a single group (Fig. 5A, Dataset 1). Most samples shared *Gluconobacter cerinus*, which was present in host fruits (see Fig. S13A in supplemental material) but absent in lab flies before egg-laying (Mothers). Among the seven ASVs classified in the second ASV group, three ASVs were present in both lab flies and other types of samples (mothers post-oviposition, G1 larvae, G1 pupae and G1 adults). Conversely, fungal samples were classified in five groups (Fig. 5B), largely corresponding to fruit types (see Fig. S13B in supplemental material). The large majority of blackberry-related samples were classified in a single group which predominantly harboured *Metschnikowia*, whereas cherry- and strawberry-related samples were classified in groups associated with *Pichia* and *Hanseniaspora* (Fig. 5B).

**Figure 5:**
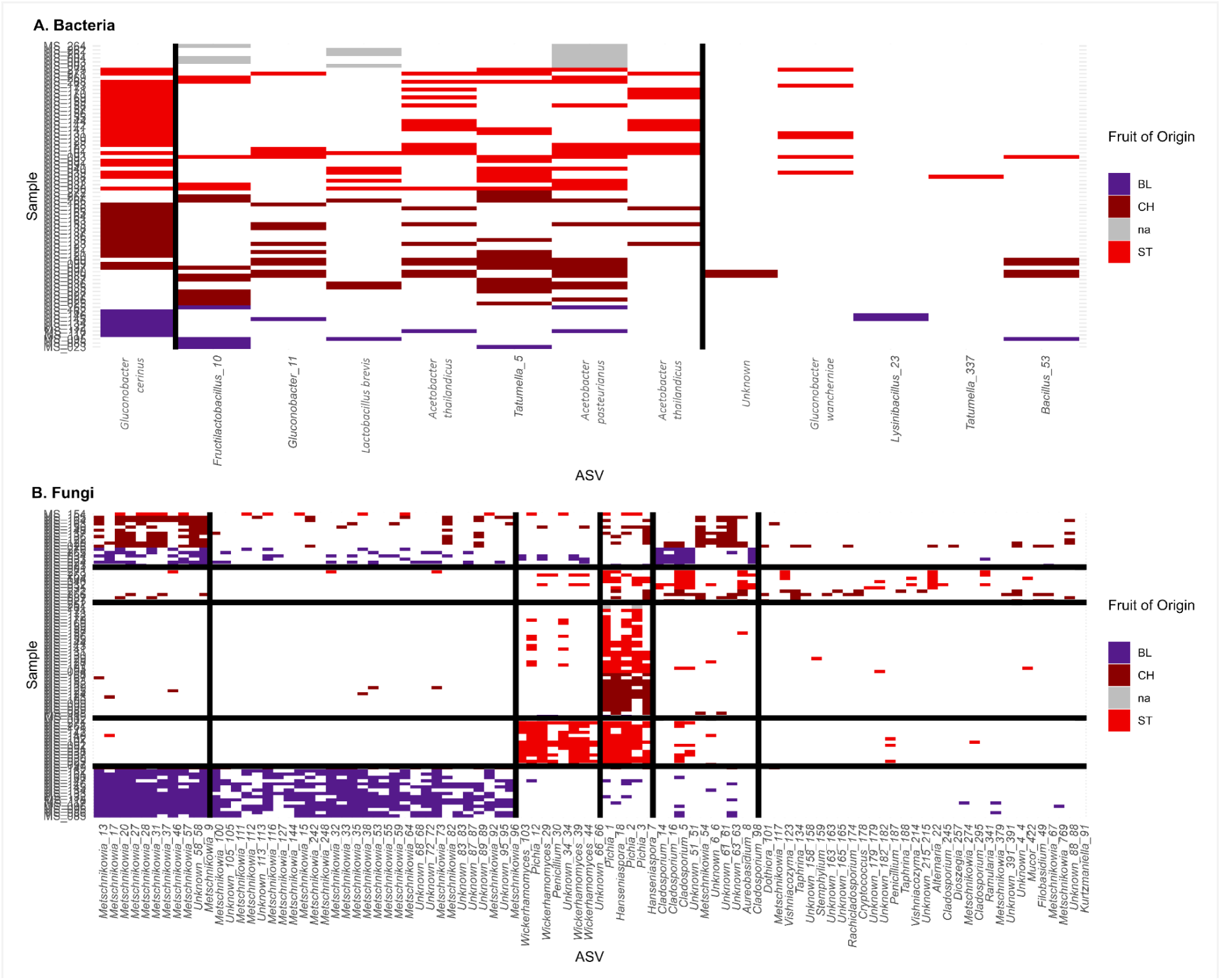
Classification of groups of *D. suzukii* samples from different fruits and across different life stages based on their bacterial (A) or fungal (B) microbial communities (presence data colored by fruit type: BL = blackberry, CH = cherry, ST = strawberry). Black solid lines delimit clusters identified based on the best latent block model. Samples with “Fruit of Origin” labelled as “NA” correspond to lab flies before egg-laying, which were not in contact with any fruits. Based on 16S (A), bacterial samples were classified into a single group (no horizontal lines divided samples in groups). Based on ITS (B), fungal samples were classified into five horizontal groups, which partially overlapped with the different fruits (bottom group included exclusively samples from blackberry, and the one above exclusively samples from strawberry).

#### Congruence of classifications

The Normalized Mutual Information (NMI) index revealed that no group defined based on LBM was specifically associated with a unique fruit, a unique life stage or sampling site and vice versa (congruence no greater than expected by chance; *p*-value > 0.05, Table 1; Fig. S14). Likewise, we did not detect any specific association between groups of fruits (Dataset 2) defined based on their bacterial or fungal communities and the type of fruit, sample type or sampling site (see Table S4 in supplemental material).

**Table 1:** Congruence between classifications of *D. suzukii* samples based on fungi presence-absence data, and potential determinants of community structure (host fruit type or sample type or host fruit sampling site), based on 10000 random permutations.

| Marker | Factor | NMI | $p$-value |
| --- | --- | --- | --- |
| ITS | Fruit Type | 0.358 | 0.1824 |
| ITS | Sample Type | 0.242 | 0.154 |
| ITS | Fruit Sampling Site | 0.499 | 0.1455 |

#### Canonical correspondence analyses (CCA)

Fungal communities were structured among host fruits and *D. suzukii* life stage (row-permutation *p*-values < 0.05, Table 2). However, this association became non-significant after controlling for differences in species richness across fruits or life stages and differences in prevalence across fungal taxa (edge-permutations *p*-values > 0.05, Table 2). Interestingly, the fungal communities of infested and uninfested fruits were strongly structured among blackberry, cherry and strawberry (both row- and edge-permutation *p*-values < 0.05, Table S5 in supplemental material), whereas bacterial communities were not (row-permutation *p*-values = 0.26, Table S5 in supplemental material).

**Table 2:** CCA between groups based on fungal communities and *D. suzukii* individuals’ life stage and their type of host fruit. *p*-value (row perm.): probability that simulated random community matrices have a *F*-statistic equal or larger to the one observed, after removing the effect of conditioning variables; *p*-value (edge perm.): probability that random community matrices, simulated while keeping the number of links per node constant, have an *F*-statistic equal or larger to the one observed. Calculations were done with 10000 random permutations.

| Formulas | ITS |  |  |  |
| --- | --- | --- | --- | --- |
|  | Chi<sup>2</sup> | F-statistic | p-value (row perm.) | p-value (edge perm.) |
| Fruit Type | 0.858 | 11.88 | 0.0001 | 0.9979 |
| Life Stage | 0.538 | 4.509 | 0.0001 | 0.8595 |
| Fruit Type + Life Stage | 1.438 | 9.768 | 0.0001 | 0.9955 |
| Fruit Type + Condition(Life Stage) | 0.900 | 14.75 | 0.0001 | 0.9978 |
| Life Stage + Condition(Fruit Type) | 0.580 | 6.414 | 0.0001 | 0.8847 |

## DISCUSSION

Our study provides an integrative characterization of the bacterial and fungal communities associated with *D. suzukii* across life stages and host fruits. Our results revealed a striking asymmetry in microbiota assembly: fungal communities are strongly structured by host fruit, whereas bacterial communities are largely shared across fruits. In contrast, both bacterial and fungal communities are influenced by developmental stages. Overall, *D. suzukii* harbours a persistent core bacterial microbiota alongside a flexible, fruit-specific, yeast community, with no evidence for strictly stage-specific microbiota.

### Distinct ecological drivers of bacterial and fungal communities of fruits

Bacterial communities associated with uninfested fruits were largely shared across fruits whereas fungal communities (yeasts) tended to be more fruit-specific (Fig. 6). The generalist bacteria *Tatumella* sp. and *G. cerinus* dominated bacterial communities across cherry, strawberry and blackberry, forming a shared core microbiota. In contrast, fungal communities comprised generalist filamentous fungi (*e.g.*, *Cladosporium*) constituting the core fungal microbiota, with fruit-specific yeasts: *Hanseniaspora* spp. and *Pichia* spp. in cherry and strawberry and *Metschnikowia* spp. mainly in blackberry. To our knowledge, few studies have studied jointly the bacterial and fungal communities across different uninfested fruits. Gurung *et al.* (35) failed to find common bacteria across their different fruit samples, including blackberry, cherry and strawberry (a review of studies on fruit microbiota is provided in Table S6 in supplemental material). The two most prevalent bacterial taxa in our study, *Tatumella* sp. and *G. cerinus*, are known to be plant pathogens (36–38), but are not commonly described as associated with the three fruits studied here. Although Zhimo *et al.* (39) identified both *Tatumella* sp. and *Gluconobacter* sp. on strawberry, both taxa (including *G. cerinus*) have never been identified together on cherry or blackberry (40–42).

**Figure 6:**
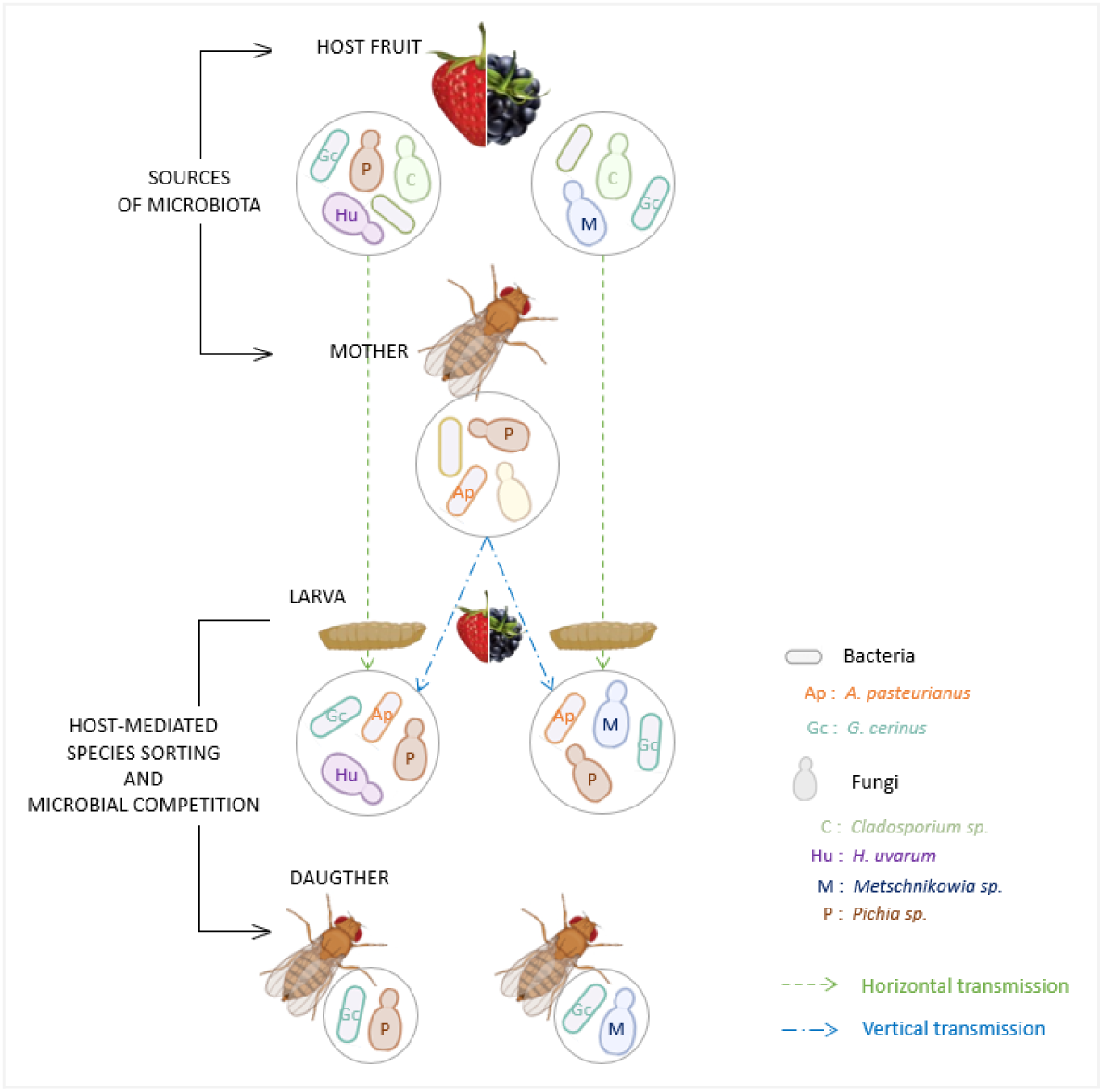
*D. suzukii* harbours a core bacterial microbiota composed of *G. cerinus*, and a niche-specific yeast microbiota transmitted horizontally from the host plant. The diversity of both bacterial and fungal communities decreases throughout development including metamorphosis.

Contrary to bacterial communities, fruit fungal communities have been more frequently studied. Consistent with our results, *Cladosporium* spp. have a wide range of host plants (43) and constitute core species found across different fruits, including blackberry, cherry and strawberry (42,43,45–51; see Table S6 in supplemental material). Also consistent with our results, *Hanseniaspora* spp. and *Pichia* spp. have been found on cherry and strawberry but not on blackberry (40,43,48,49; see Table S6 in supplemental material). In contrast, studies have found *Metschnikowia* spp. on cherry and strawberry (42,47,48) or on blackberry (51). Interestingly, our results suggest that *Metschnikowia* sp. (likely *M. pulcherrima*) might be structured between a taxon found across all three fruits and a more blackberry-specific taxon. Further studies will shed light on this potentially hidden genetic structure.

We observed an asymmetric distribution pattern between bacteria and filamentous fungi vs. yeasts on fruit surfaces (Fig. 6). These differences could be due to neutral or selective processes, both of which are not mutually exclusive. Indeed, yeasts could have smaller dispersal rates and/or stronger ecological filtering than bacteria and filamentous fungi. Our understanding of the mechanisms that govern the structure of microbial communities on fruit surfaces remain limited, as studies have focused more on temporal changes (*e.g.* over ripening stages (52) or seasons (49)). The asymmetric spatial repartition of bacterial and filamentous fungi vs. yeasts we observed could be explained by heterogeneous movements and dispersal ability (53). While bacteria and filamentous fungi are capable of both wind-driven and insect-mediated dispersal (*e.g.*, via ascospores (54–56)), yeasts are primarily dispersed by insects (57–59). Differences among insect vectors in dispersal toward different fruits could result in the observed fruit-specific distribution. Although insects pollinating blackberries, cherries and strawberries are similar, including common bee species (60–62), attraction to different fruits could vary among species.

Second to dispersal, ecological filtering by fruit surface environments could additionally or alternatively explains the observed asymmetric spatial repartition. Both bacteria and filamentous fungi often use a wide range of plant-derived compounds compared to yeasts. Bacteria are indeed very efficient at metabolizing organic substrate (63) and filamentous fungi produce a broader array of secondary metabolites compared to unicellular yeasts (64), which frequently show niche-specialized metabolic repertoires (65). Moreover, niche partitioning is an important determinant of fungal community structure (66). Local specialisation of yeasts, and their potential dispersal limitation compared to bacteria and filamentous fungi, as mentioned above, could then explain yeast fruit-specificity. Such a pattern is consistent with theoretical predictions that specialisation is promoted by a limited dispersal (53,67). Together, our results suggest that microbiota assembly on fruit surfaces might arise from both dispersal limitation and ecological filtering, with differing contributions across microbial groups.

### Core bacterial vs. niche-specific fungal microbiota in *D. suzukii*

We identified *G. cerinus* as a core bacterial taxon of *D. suzukii* microbiota, consistently present across all life stages for cherry or strawberry, and across pupae and emerging adults for blackberry (Fig. 6). In addition, this taxon was absent in lab flies but present in uninfested host fruits and females after oviposition on fruits. This pattern is consistent with the horizontal acquisition of some bacteria from each host fruit by *D. suzukii* larvae followed by the specific host filtering of *G. cerinus*, with preservation throughout development and metamorphosis. Few studies characterized the microbiota associated with *D. suzukii* larvae across different host fruits (see Table S7 in supplemental material for a synthetic review). Associations between *D. suzukii* and *G. cerinus* have been documented in larvae (31) or field-caught adults (68,69). These results suggest a stable association despite environmental acquisition. *G. cerinus* might be an ecologically and functionally important microbial species for this fly pest due to its ability to metabolize major amino acids (70). In addition, He *et al.* (37), found beneficial effects of *G. cerinus* on the growth of *Bactrocera dorsalis* larvae. Further experiments with field-caught females would be needed to verify the transmission patterns we observed, and to identify whether *G. cerinus* is passively acquired and maintained, or actively selected, by the host because of beneficial effects on its performance. In addition, other bacteria showed more variable transmission patterns either from the fruit (*Acetobacter thailandicus, Tatumella* spp.) or from the mothers to their offspring (*A. pasteurianus*). Consistently with our results, these species have been previously identified as commonly present in *D. suzukii* microbiota in the wild (31,35,68,71). Pais *et al*. (5) also showed that *A. thailandicus* formed a stable beneficial association with *D. melanogaster* in the laboratory. These bacteria might form more flexible and transient associations with *D. suzukii*.

Contrary to bacteria, no core fungal microbiota was detected. We observed that dominant yeast taxa associated with *D. suzukii* differed among fruit types (*Metschnikowia* spp. in blackberries vs. *Hanseniaspora* spp. and *Pichia* spp. in cherries and strawberries, Fig. 6). The different permutation-based tests show that fungal communities structure among fruits was due to differences in richness between blackberry vs. cherry and strawberry. The observed NMI between LBM sample clusters and fruit type did not significantly exceed the values obtained from Curveball-simulated networks, indicating that the observed correspondence was not stronger than expected under the null model (Table 1; Fig. S14 in supplemental material). Furthermore, the CCA with row permutations revealed a significant association between fruit type and the sample classification identified by the LBM, based on sample fungal communities. However, the association was not significant under the Curveball-based configuration null model, which preserves sample richness and taxon prevalence (edge permutations, Table 2). This result suggests that significant row-permutations may be due to differences in species richness or prevalence among samples rather than consistent differences in community composition among fruits. This interpretation is consistent with our observation that fungal species richness was higher in individuals from blackberries than those associated with cherries or strawberries (Fig. 3B).

In addition, *Metschnikowia* spp. and *Hanseniaspora* spp. were absent in lab flies but present in uninfested host fruits and females after oviposition on fruits. This observation is consistent with the horizontal acquisition of yeasts by *D. suzukii* larvae from their respective host fruits, followed by host-mediated filtering of some taxa which persist throughout development. Those two genera and *Pichia* spp. are commonly found in *D. suzukii* (72,73) and their host fruits (47,74,75). While *Hanseniaspora* has been described as being greatly attractive for *D. suzukii* and supporting larval growth (47,76), *Pichia* and *Metschnikowia* have been described as harmful for the survival of larvae (77,78) and pupae (8) - although *Pichia* has also been described as beneficial for *D. suzukii* (4,76). Whether fruit-specific yeasts are beneficial for larval development on different fruits remains to be investigated.

Furthermore, we observed the transmission of a *Pichia* sp. from females to their offspring across fruits (Fig. 5B), suggesting that the species was transmitted throughout metamorphosis (also observed for *A. pasteurianus*; Fig. 6). A pseudovertical transmission could explain these observations (25). Indeed, as mothers and their offspring partially share the same environment (host fruits), substrate inoculation with microbes species could allow for the transmission (5). Although we verified that the more abundant bacterial and fungal families composing *D. suzukii* larval microbiota in the lab and from the field were similar, whether the transmission of *Pichia* spp. and *A. pasteurianus* occurs in natural populations remains to be investigated.

In other holometabolous polyphagous insects, microbial associations are mostly depicted as not persistent through life stages (79) and generations (80), against the generalisation of a core microbiota. Yet, the host plant is also commonly described as the main driver of both fungal and bacterial microbiota composition (79–82). Besides, the acquisition and horizontal transmission of facultative microbes through host plants are processes which are conserved among Holometabola, although mostly demonstrated with bacteria (78) and being species-dependent. It is therefore difficult to use our observations to suggest a general pattern regarding the microbiota associated with holometabolous polyphagous insects, other than highlighting the importance of the host plant.

### Selection reduces microbial diversity during *D. suzukii* metamorphosis

Both bacterial and fungal diversity declined during *D. suzukii* development, consistent with patterns observed in other holometabolous insects. The transient period when emerging adults harbour depleted communities is likely shortened in the wild, as insects in nature are directly in contact with multiple microbial sources to replenish their associated communities and as adults usually harbour a more diverse microbiota in the wild than in the laboratory (81). For instance in certain lepidoptera, bacterial richness declines sharply between larval and pupal stage in the wild, followed by partial recovery in adults (82). In contrast, in tephritids flies a strong bacterial diversity reduction occurs post-pupal emergence in laboratory populations (83), but in the wild diversity is conserved over life cycle (84,85). Thus, additionally to interspecific variations, environmental conditions (natural vs. laboratory conditions) are important factors to consider when analyzing microbial communities over insects’ life cycle.

Bacterial communities shifted towards Acetobacteraceae in pupae and emerging adults at the expense of Lactobacillaceae and Erwiniaceae. This contrasts with the patterns of a drastic increase of *L. fructivorans* from pupae to young adults described in a lab strain of *D. melanogaster* (27), highlighting possible species-specific dynamics. In emerging adults, *Pichia* and/or *Metschnikowia* dominated over *Hanseniaspora* and *Cladosporium*, consistent with results in *D. melanogaster* (8,86). Population dynamics, such as competition for substrate nutrients likely occurred between species on fruit substrate. In particular, as previously described on apple surface (87), *Metschnikowia* is likely to outcompete *Pichia* on fruit surfaces, as we found that *Pichia* was associated with the absence of *Metschnikowia*. The compositional shift we observed in *D. suzukii* might be host-specific without being generalisable to other holometabolous polyphagous insects. Indeed, the bacterial or yeast genera we identified in emerging adults were not found in other fruit flies species (27,84,85).

### Potential limitations of our study

Our failure to detect fruit-specific bacterial taxa could be due to the filtration threshold of 60 % prevalence across samples. To verify this, we performed the analyses after filtering ASVs in Dataset 1 using a less stringent threshold of 20 % prevalence across samples. We observed the same trend between rarefaction curves using 60 % and 20 % thresholds (Fig. S3 vs. S4 in supplemental material), and we did not observe the apparition of fruit-specific taxa when performing ordination after using a relaxed threshold (Fig. 4 vs Fig. S12 in supplemental material). Thus, the uniformity of bacterial communities across samples from different host fruits was not an artifact of our stringent filtering.

Our results regarding the horizontal transmission and host filtering of *G. cerinus*, *Metschnikowia* spp. and *Hanseniaspora* spp. appear robust. However, the potential pseudovertical transmission of *Pichia* spp. or *A. pasteurianus* observed from laboratory mothers to their offspring might be an artifact of laboratory adaptation. Indeed, the artificial medium used to maintain our laboratory populations of *D. suzukii* is almost sterile, so that selection for pseudovertical transmission of the microbiota is very strong in the laboratory: in the absence of microbial transmission larvae cannot develop on artificial medium. Further studies with field-collected females infected with *Pichia* spp. or *A. pasteurianus* will be necessary to validate this pseudovertical transmission.

Our controls showed that the microbiota of larvae naturally infested in the field was similar to larvae obtained through artificial infestation in the lab. Besides, after the transfer of these larvae to artificial medium, changes in microbial communities in pupae and adults were similar independently of whether the larvae originated from natural or artificial infestation. However, in nature horizontal microbial transfer from decaying fruit to pupae and emerging adults may occur as these stages complete their development on fruit (35,88), a process that could not be captured under our experimental conditions. Further studies will show whether pupae and newly emerged adults in the field also show a strong reduction in the diversity of bacterial and fungal taxa.

### Conclusion

Our results show that fruit-associated bacterial and fungal communities are shaped by distinct ecological processes. Bacteria and filamentous fungi appear widely distributed, whereas yeasts are more strongly structured among host fruits.

In *D. suzukii*, this translates into a core bacterial microbiota shared across life stages and fruits, and a niche-specific yeast microbiota structured by host fruits. These findings suggest that yeast and bacteria might play complementary roles in insect ecology, and emphasize the need to study them jointly. Taken together our results underline how using a metacommunity approach, by focusing on ecological processes driving communities assembly (e.g. microbial dispersal among fruits, horizontal transmission and host filtering across life stages), may provide a deeper understanding of tripartite interactions among insects, host plants, and their microbiota.

## MATERIAL AND METHODS

### Fly populations and rearing

We used two long-term laboratory populations of *D. suzukii* initiated from a field sampling and maintained for more than 100 generations on artificial media composed of either cherry or strawberry puree (89). During the experiment, flies were reared under constant conditions: 21 °C and 60 % relative humidity with a 16:8 light:dark cycle. Because flies have been maintained under stable conditions for many generations, we assumed that they harbour relatively stable microbial communities, potentially reduced in diversity compared to wild populations (4,35).

### Fruit sampling

In June 2023, cherries and strawberries were collected from two sites, while blackberries were collected from one site in southern France (see Fig. S1 and Table S1 in supplemental material). Except for the blackberries which were bought, fruits were collected in the field, brought to the laboratory the day before starting the experiment and kept overnight at 5 °C (Fig. 7, step 1).

**Figure 7:**
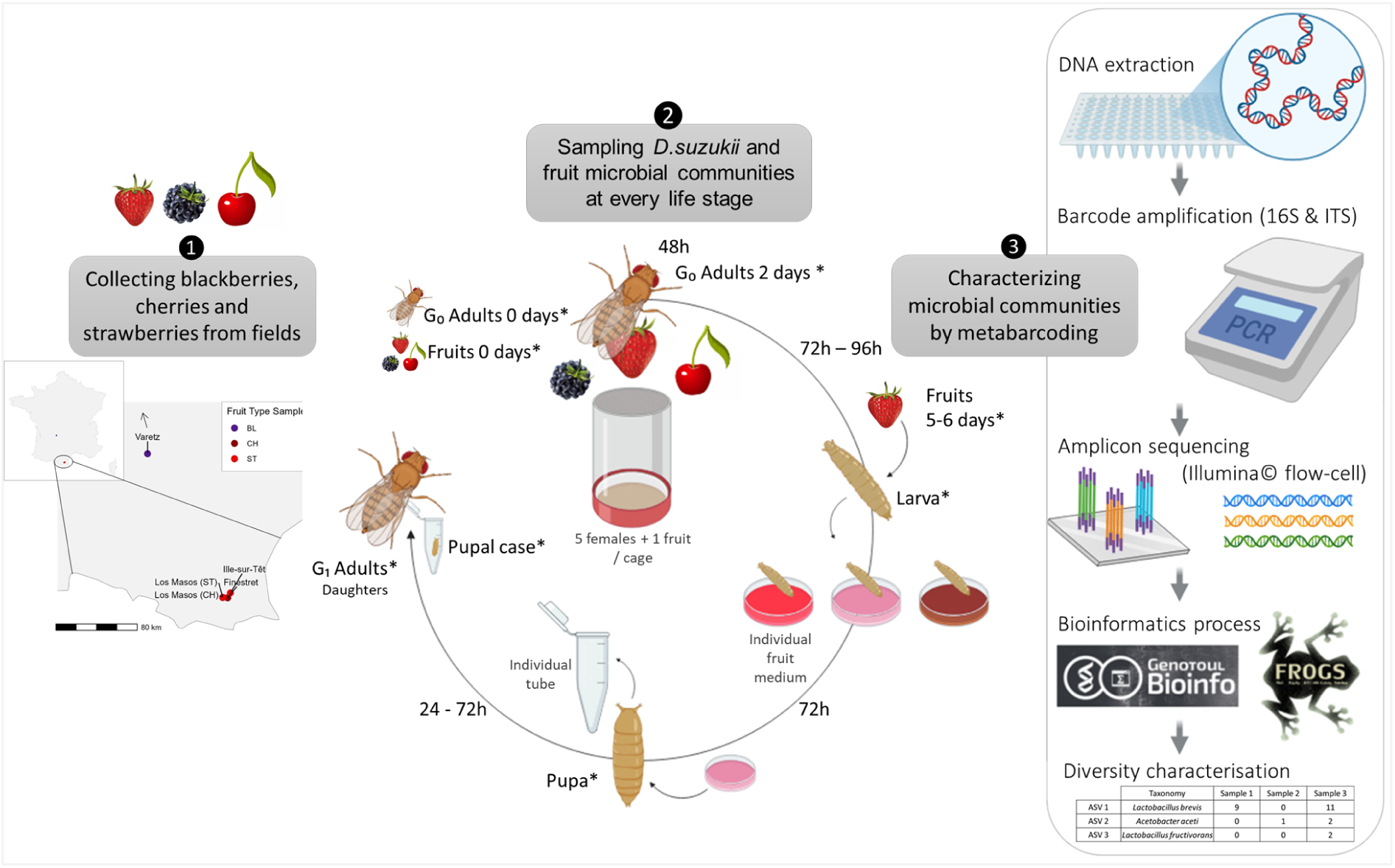
Experimental design to test for the effect of host fruit type and fly life stage of *D. suzukii* microbiota composition. * stars indicate samples that were frozen at −70 °C for 16S and ITS metabarcoding.

### Experimental procedure

To compare the compositions of the microbial communities of individuals across life stages and among host fruits, we sampled egg-laying mothers, larvae, pupae, pupal cases and emerging adults from all host fruits (Fig. 7, step 2). As controls, we also included naturally infested fruits collected from all sites except one (no infested blackberry could be found), larvae extracted from these fruits, and pupae and emerging adults that developed from these larvae on fruit-based media (see Doc. 1 in supplemental material).

#### Step1.1 Host fruits and flies sampling

At the start of the experiment, subsets of fruits were frozen in individual sterile plastic bags (12.5 x 12.5 cm, Whirl-Pak, FisherScientific). Later, fruits were thawed and juice was extracted inside the sterile bags for 16S and ITS metabarcoding (“Fruits 0 days”, 16 to 25 pools for each of the five types of fruits, Fig. 7). To characterize maternal microbiota prior to oviposition, five pools of five mated females were frozen in sterile tubes (2-mL, Eppendorf) for metabarcoding (“G_0_ Adults 0 days”, Fig. 7).

#### Step1.2 Fruit artificial infestation and adult females sampling

To artificially infest fruits with larvae, we anaesthetized flies and manually separated females from males. For each sampling site, 10 fruits were individually exposed to five mated females in an embryo collection cage (3.75 (D) x 5.8 cm (H), Genesee Scientific) for 48 hours to allow oviposition. Experimental controls were also implemented, particularly to confirm that natural infestation could be neglected relative to artificial infestation (see Doc. 1 in supplemental material). After 48 hours, we removed females from the cages after putting them on CO₂ pads. Live females were distinguished from dead ones and frozen in sterile tubes for metabarcoding (“G₀ Adults 2 days”, 12 to 20 pools per fruit type, Fig. 7).

#### Step1.3 Larvae extraction from host fruits

Seventy-two hours after females removal, we extracted G₁ larvae by removing the stem or calyx from fruits and pressing them individually in a sterile plastic bag. Larvae were rinsed with sterile PBS before being individually transferred into a 96-well plate with artificial fruit medium to develop. To avoid bias associated with a single fruit medium, larvae from each fruit were evenly distributed between blackberry-, cherry-, and strawberry-based artificial media. To prevent larval exposure to other microbes (than from mothers and fruits in which they hatched) artificial fruit media were previously pasteurized (15 minutes at 60 °C). To prevent airborne contaminations and larvae escape, plates were sealed with a sterile and breathable adhesive film (AeraSeal adhesive film, Excel Scientific, Wrightwood, USA).

#### Step1.4 Larvae and host fruits sampling

To compare microbial communities of G₁ larvae with the community of their mothers and host fruits, we collected additional larvae from each host fruit, froze them in sterile tubes and later crushed them to extract DNA for metabarcoding (“Larvae”, 8 to 13 pools per host fruit, Fig. 7). To investigate the effect of larval infestation on fruit microbiota, we also sampled the juice of fruits from which larvae were extracted, froze it in sterile tubes for metabarcoding (“Fruits 5-6 days”, 5 to 11 pools per host fruit, Fig. 7).

#### Step1.5 Pupae extraction from fruit media and sampling

Three days after larvae transfer, the presence of G₁ pupae in wells was controlled every two days, except on weekends. The 96-well plates were checked with a magnifier under a laminar flow cabinet. Pupae were extracted from fruit media and transferred individually in a sterile tube. To characterize pupae microbial community, we froze every other tube with a pupa for metabarcoding (“Pupae”, 11 to 21 pools per host fruit, Fig. 7).

#### Step1.6 Adults emergence, pupal cases and emerging adults sampling

From the first pupa appearance, emergence of G₁ adults was controlled every day, except on weekends. Emerging flies were separated from their pupal cases (puparium), and frozen separately for metabarcoding (“PupalCase”, 4 pools per fruit type; “Daughters”, 12 to 24 pools per fruit type, Fig. 7).

### Samples preparation, DNA extraction, 16S and ITS metabarcoding

We prepared more than 450 samples for metabarcoding by pooling five biological replicates (individuals or fruit juice), or fewer if not available (min. two). To detect only the internal microbiota of individuals (except pupal cases), pools were surface-sterilized through three 30-second baths of bleach at 50 %, followed by two baths of sterile water (90), before being transferred with sterile forceps in microtubes for DNA extraction.

DNA was extracted and purified using DNeasy® 96 Blood & Tissue kit (Qiagen), following the protocol of Chapuis et al. (91) (see Doc. 2 in supplemental material for details). The V4 region of 16S rRNA gene was amplified by PCR (Fig. 7, step 3) using custom-made indexed primers (92), and the ITS1 region of ITS rRNA using BITS and B58S3 primers, as previously described (93). We used ZymoBIOMICS microbial community DNA standard (Zymo Research Corp.) and Mycobiome Genomic DNA Mix (ATCC, MSA-1010) for 16S and ITS1 positive DNA controls. To detect contaminations from the DNA extraction kit or the PCR mix, we included negative extraction and PCR controls (no sample or no DNA respectively). To control for index switching during sequencing, empty wells corresponding to unused barcodes combinations were included (92). We pooled all PCR amplicons for 16S and ITS separately, purified pooled DNA using a gel-extraction kit (NucleoSpin Gel and PCR Clean-up, Macherey-Nagel) and quantified them by qPCR (kit KAPA SYBR® FAST qPCR, AffiPCR Biosystems). To calibrate the sequencing depth of each type of sample for both markers, we used independent preliminary data from another study (see Table S2 in supplemental material). Normalized amplicon libraries were sequenced at Montpellier Genomics and Sequencing Platform (GenSeq, MEEB, University of Montpellier) using an Illumina MiSeq Reagent Kit v2 (2 x 250 cycles). Samples with initial low coverage were sequenced twice or three times (171 and 48 samples for 16S and ITS respectively) using two additional sequencing runs.

### Bioinformatic analyses

All analyses were performed using R (version 4.3.1; R Core Team 2023). We performed bioinformatic analysis following Chapuis *et al.* (91). Briefly, demultiplexed paired-end reads were trimmed with Cutadapt v1.8.3 (94) and merged into contigs with FLASH v1.2.6 (91,95). For the rest of bioinformatic analyses, we used the FROGS 4.1 pipeline (96). We excluded contigs that did not match the range of expected length (16S: 100-241 bp; ITS: 50-700 bp). To dereplicate sequences, we clustered them into Amplicon Sequence Variants (ASVs) using Swarm v2 (parameter d=1; (97)). We removed PCR chimeras in each sample and filtered out false positives that arose during the PCR library preparation or through index switching during sequencing. To correct for random contamination, we removed ASVs when they occurred in only one of the two PCR replicates. Taxonomy of 16S and ITS ASVs were assigned using SILVA 138.2 pintail100 (98) and UNITE Fungi 10.0 (99) respectively.

Taxonomic and abundance tables were cleaned using R scripts (91) with minor modifications: multi-affiliated ASVs assigned to a specific taxon more than 50 % of the time were assigned to this taxon. We removed potential undetected residual chimeric contigs using the function isBimeraDeNovo from dada2 package (100). The taxonomy of undetermined ASVs within the 1000 most abundant ASVs were manually found by BLAST in the NCBI database, which represented 20 bacterial and 78 fungal ASVs. The ASVs corresponding to plants, chloroplasts, mitochondria, or Fungi kingdom (respectively Bacteria) were removed from 16S abundance table (resp. from ITS). We manually removed ASVs corresponding to *Wolbachia* (43 % of 16S reads) because it is an intracellular vertically transmitted endosymbiont and it can dominate sequencing reads without reflecting environmentally acquired microbiota. We also manually removed likely contaminations from human microbiota which represented 107 bacterial and 30 fungal ASVs (see Table S3 in supplemental material), ASVs present in no sample, and samples with no reads. This filtering resulted in 249 16S and 288 ITS samples remaining over the 298 of interest. The distribution of reads per sample before and after those filtering steps is reported on Fig. S2, in the supplemental material.

We then divided our data in three datasets for later analyses. Dataset 1 included *D. suzukii* individuals (mothers before and after egg-laying, larvae, pupae and emerging adults) and their host fruits, before and after infestation. Dataset 2 included only host fruits (uninfested or naturally infested) with two and four supplementary sampling sites respectively for blackberry, and cherry and strawberry (see Table S1 in supplemental material). Dataset 3 included *D. suzukii* individuals (larvae, pupae and emerging adults) from lab populations used for artificial infestations and from wild populations present in fruits naturally infested. To filter rare or sample-specific ASVs, we calculated the prevalence of each ASV across all combinations of life stages and host fruits for Dataset 1 and Dataset 3 (or fruit state and host fruit for Dataset 2), and removed ASVs that failed to reach a prevalence of 60 % in at least one combination, excluding groups with less than two samples (except for Dataset 3 bacterial ASVs which were filtered above 50 % of prevalence). It excluded 93.1 % of bacterial and 75 % of fungal ASVs from Dataset 1, representing only 6.91 % and 7.21 % of total reads. For Dataset 2, 96.2 % of bacterial and 84.5 % of fungal ASVs were removed, representing respectively 11.5% and 8.82 % of total reads. For Dataset 3, 89 % of bacterial and 71.9 % of fungal ASVs were removed, representing respectively 10.8 % and 11.5 % of total reads.

Finally, to correct for uneven sequencing depth among samples, we rarefied depth to 3,000 sequences per sample for 16S, and 5,000 sequences per sample for ITS in both datasets. Most rarefaction curves approached saturation, indicating that our sequencing depth captured microbial diversity at large, despite the lost bacterial sequencing depth due to chloroplasts, mitochondria and *Wolbachia* (see Fig. S3 in supplemental material). We conserved 130 samples and 16 ASVs for 16S, and 163 samples and 116 ASVs for ITS in Dataset 1; respectively, we conserved 56 samples and 29 ASVs for 16S, and 89 samples and 109 ASVs for ITS in Dataset 2, and 60 samples and 10 ASVs for 16S, and 163 samples and 116 ASVs for ITS in Dataset 3.

### Statistical analyses

#### Community description

We first described the composition and diversity of bacterial and fungal communities harboured by *D. suzukii* and its host fruits. We identified the core microbiota (*i.e.* present across most host fruits and life stages), the fruit-specific microbiota (*i.e.* present across most fly life stages from a given host fruit), and the life stage-specific microbiota ASVs (*i.e.* present in a given fly life stage across most of fruits). We analysed alpha and beta diversity of communities using the Phyloseq package (101) on Dataset 1. We calculated different metrics (observed richness, Shannon and Chao1) to compute alpha diversity of microbial communities. To explore beta diversity, we used an ordination on presence/absence matrices on non-metric multidimensional scaling (NMDS) using Jaccard distance metric.

#### Hypothesis testing

We tested for differences in microbial communities of *D. suzukii* individuals depending on life stage, on host fruit or sampling site using network analyses. We assessed the congruence of sample classification obtained through community-search algorithms with those associated with our three categorical variables of interest (102,103). To search for communities within the network, we inferred sample classification using a latent block model (LBM) algorithm with ‘estimateBipartiteSBM’ function from the SBM package (R package version 0.4.7; (104)). We first assessed separately the congruence between the sample classification found with the LBM and each variable of interest (life stage, host fruit or sampling site) using the Normalized Mutual Information index (NMI). NMI measures the congruence from zero to one between two classifications. The significance of the NMI was tested by comparing the observed NMI to 10000 NMI values simulated using randomized networks (obtained with the ‘curveball’ algorithm implemented in the ‘compare’ function; igraph package v2.2.1; (105)). Second, we assessed the congruence between the classification found with the LBM and both variables of interest using a Canonical Correspondence Analysis (CCA). CCA decomposes the variation in the community found with the LBM (nodes classification) through projections into the eigen spaces induced by the two external variables tested (102). Last, we tested for the significance of the two factors’ effects using row permutations. If significant, we used a second randomization keeping the number of links per node constant, to verify whether differences in microbial communities among factor levels was still significant after accounting for differences in richness among factor levels (102).

## DATA AVAILABILITY

The datasets and R scripts used and/or analyzed in this study have been deposited in the GitHub repository https://github.com/sidodq/Y2025.TransMicro2. Raw reads are available at NCBI SRA database under BioProject PRJNA1514060.

## ACKNOWLEDGEMENTS

We thank the farmers who provided the fruits. We thank Amélie Sibrao, Maxime Galan, Svitlana Serga and Guénaëlle Genson for their technical support during larval sampling or metabarcoding. We thank Cécile Neuvéglise and Romain Gallet for helpful discussions. We contributed to the present study as follows: N.O.R was responsible for the overall conception and supervision of the study and acquired funding; N.O.R., S.D., M.L., C.D. and L.C. designed and conducted the sampling and rearing; S.D., M.L. and A.L. conducted the metabarcoding; M.L. and S.D. analyzed the data; S.D., B.F. and N.O.R. wrote and edited the manuscript; all authors read and approved the manuscript. Data presented in this work were partly produced through the GenSeq technical facilities of MEEB (CNRS and University of Montpellier) hosted by ISEM (CNRS, University of Montpellier and IRD). AI tools were punctually used for R script writing. This project received funding from the European Regional Development Fund (ERDF) and from the Agence Nationale de la Recherche (ANR-22-CE20-0002-01).

## SUPPLEMENTAL MATERIAL

### 1. Supplemental Figures

**Figure S1:**
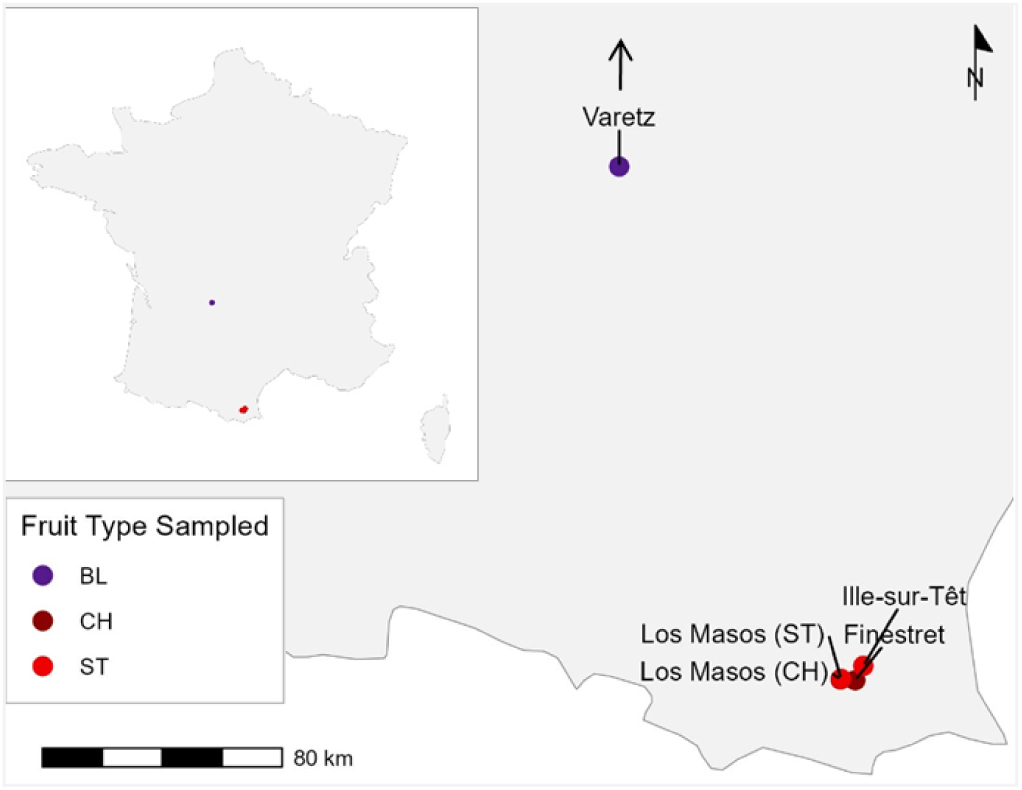
Geographic locations of the sampling sites of the different fruits for Dataset 1. BL: blackberry, CH: cherry, ST: strawberry

**Figure S2:**
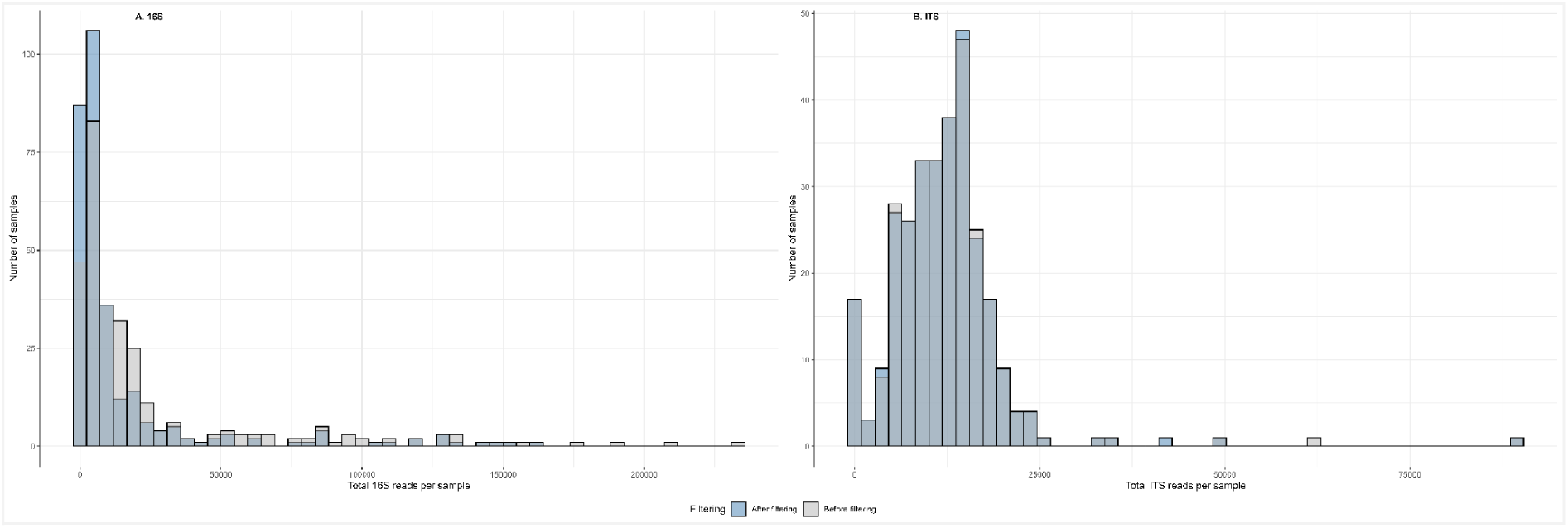
Distribution of reads per sample from Dataset 1, before and after filtering out Wolbachia, plant-derived and contaminant reads for 16S (A) and ITS (B) marker sequencing

**Figure S3:**
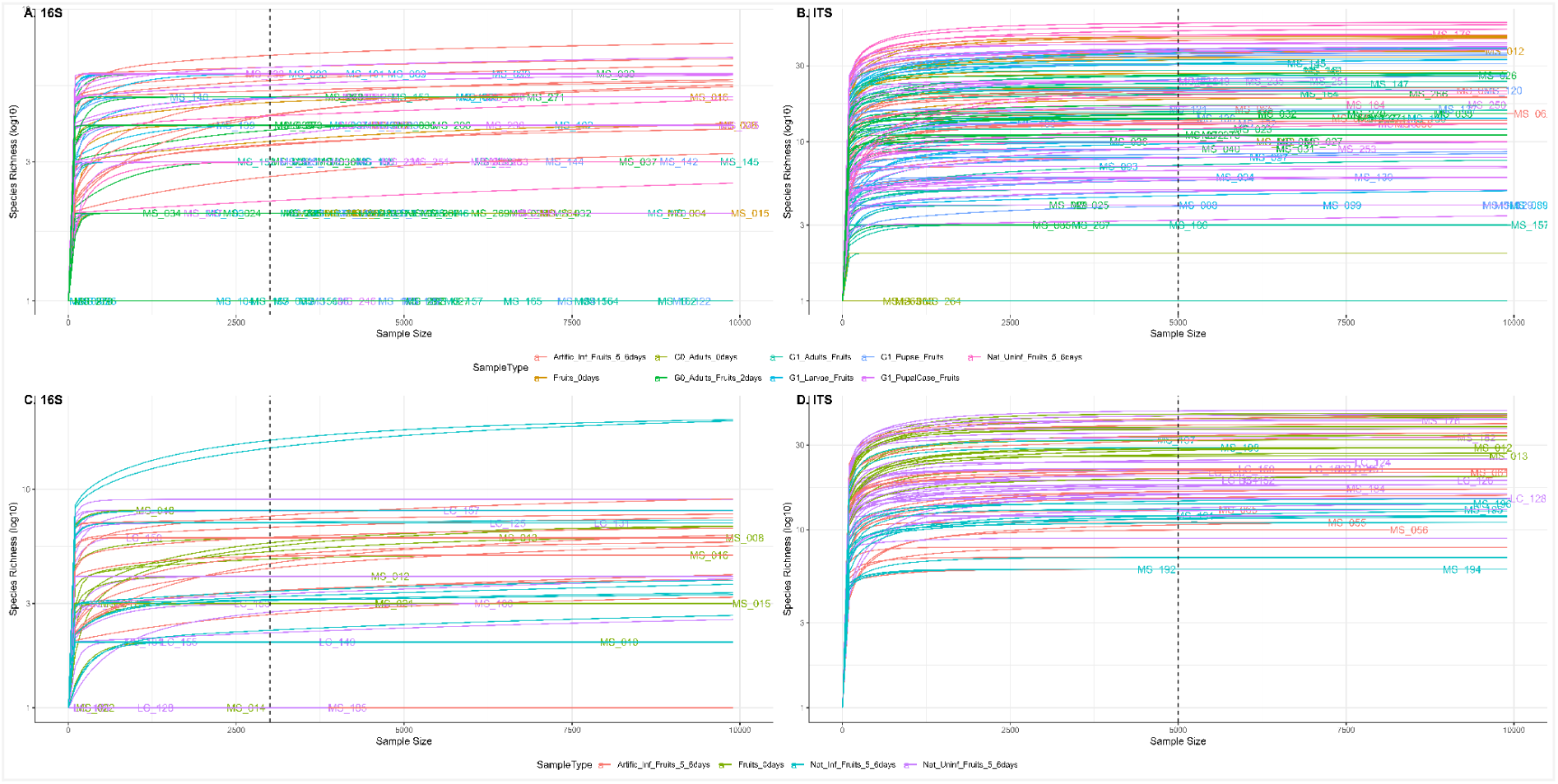
Rarefaction curves for Dataset 1 (A, B) and Dataset 2 (C, D) based on 16S (A, C) and ITS (B, D) markers after filtering ASVs with a 60% prevalence threshold across samples. Dashed vertical lines represent the rarefaction threshold of 3,000 reads for 16S and 5,000 reads for ITS sequencing data.

**Figure S4:**
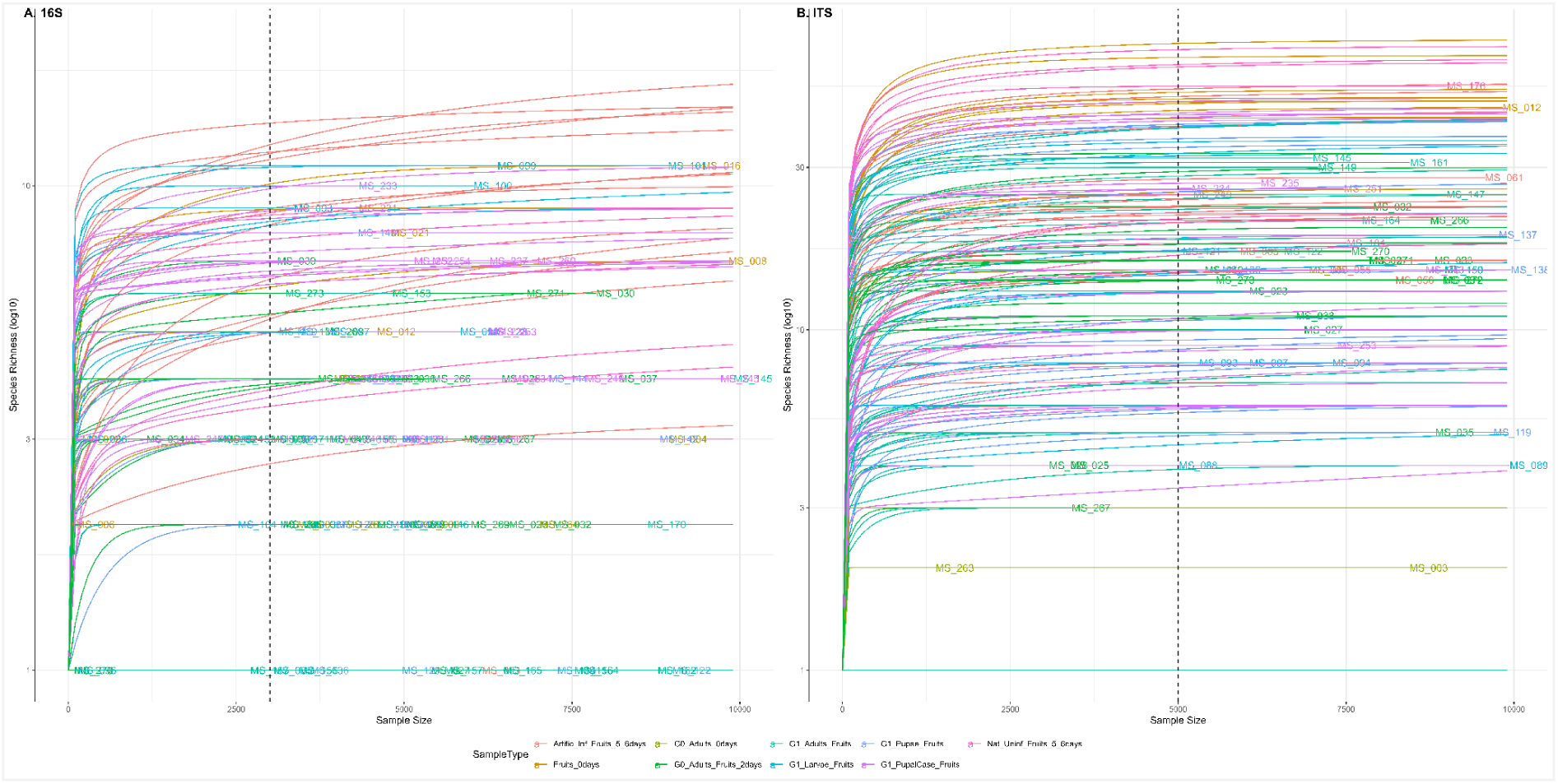
Rarefaction curves of samples from Dataset 1 sequenced using 16S (A) and ITS (B) markers after filtering with a threshold of 20 % on ASV prevalence across samples. Dashed vertical lines represent the rarefaction threshold of 3,000 reads for 16S and 5,000 reads for ITS sequencing data.

**Figure S5:**
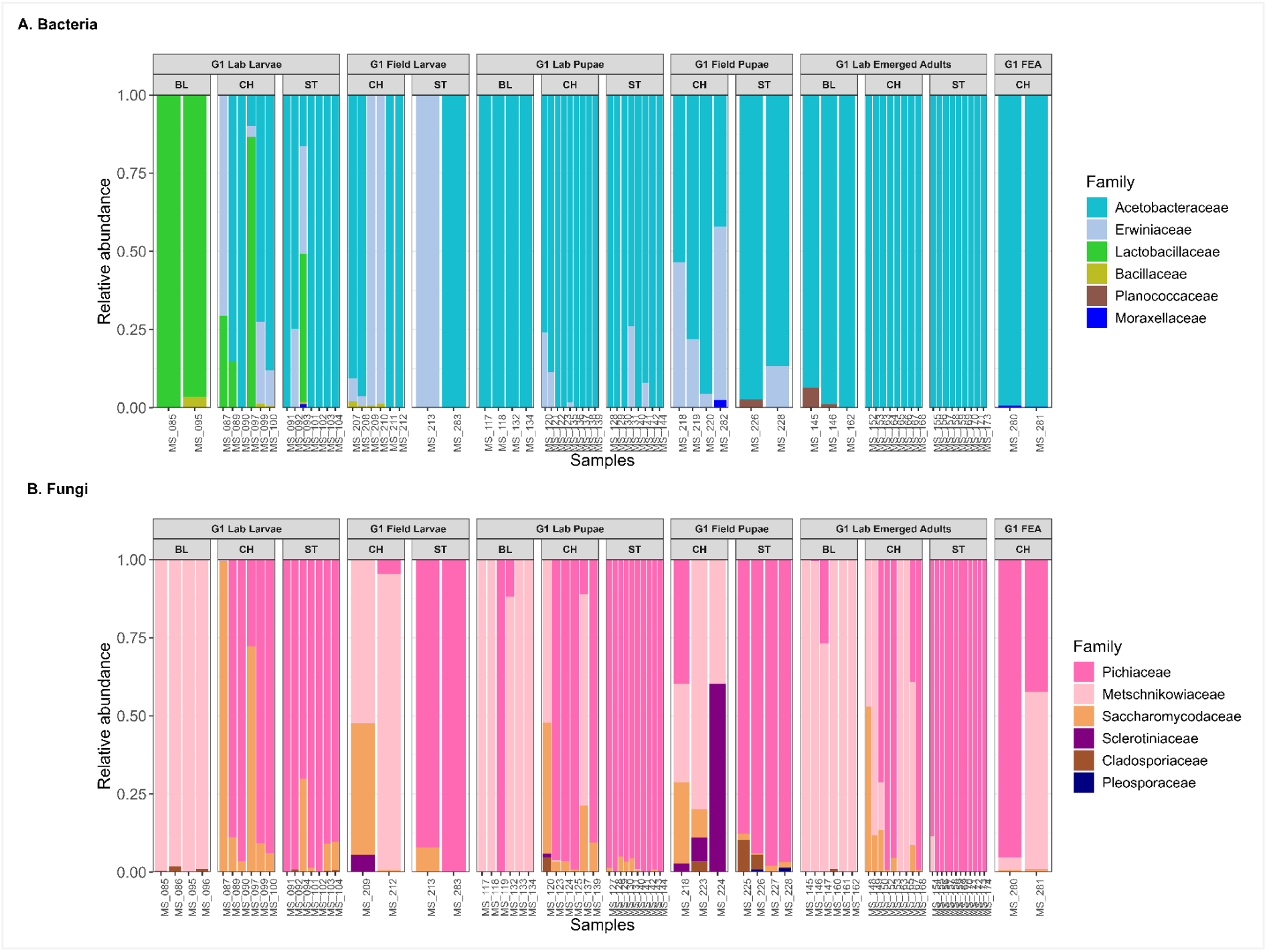
Relative abundance of the 6 more abundant bacterial (A) and fungal (B) families associated with *D. suzukii* individuals from fruits artificially infested in the lab and individuals from fruits naturally infested in the field, separated by life stage and host fruit (Dataset 3). G1 FEA: Fruits Emerged Adults, i.e. adults emerged in the lab after the transfer of larvae from naturally infested fruits’to fruit media. BL: blackberry, CH: cherry, ST: strawberry. No data were available for field individuals related to blackberries as fruits were naturally free from *D. suzukii* infestation.

**Figure S6:**
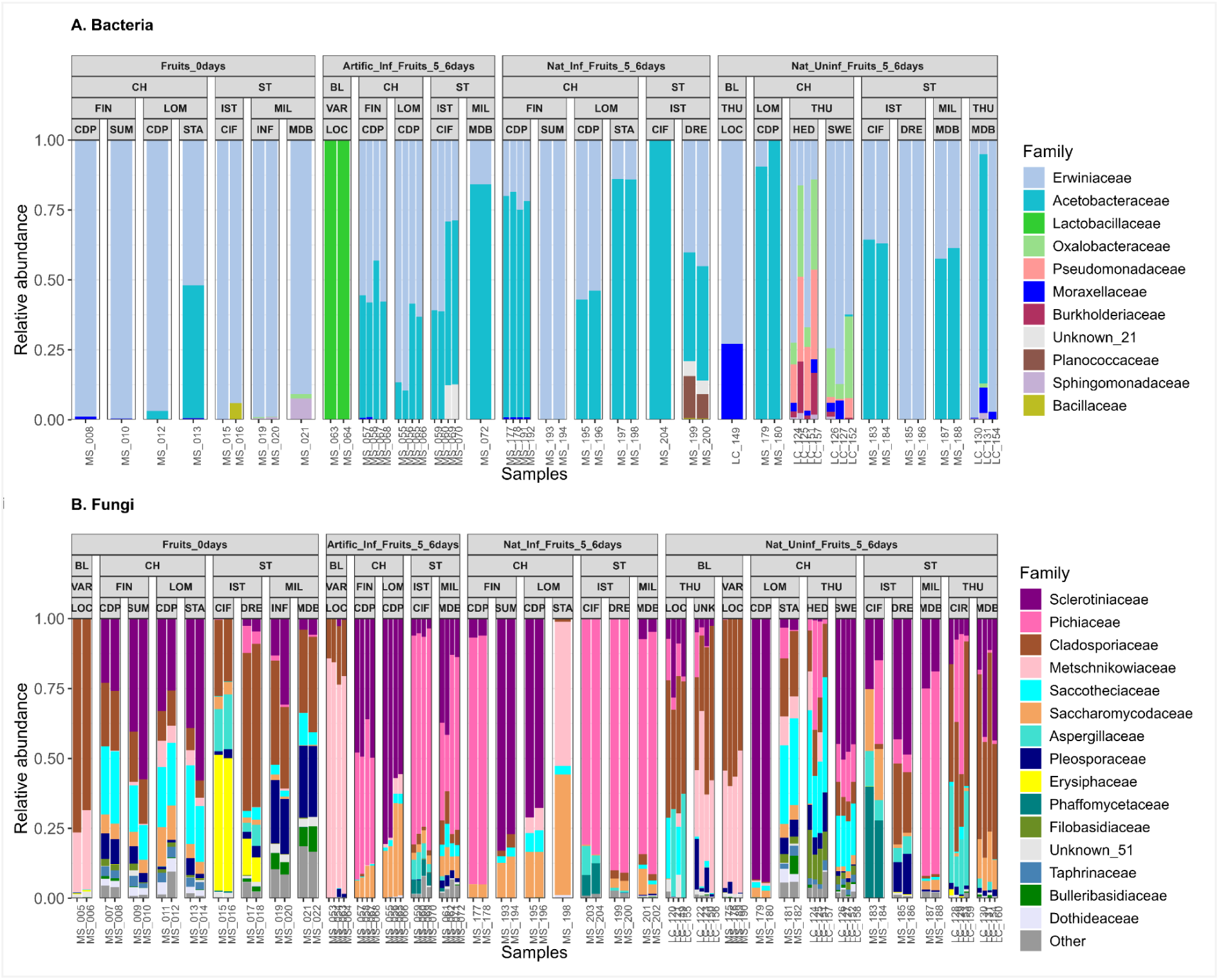
Relative abundance of the 11 more abundant bacterial (A) and 15 fungal (B) families present on infested and uninfested fruits, separated by sample type, fruit type, sampling site and fruit variety (Dataset 2). Sample type: uninfested fruits at 0 days (Fruits_0days), fruits artificially infested with *D. suzukii* in the lab at 5-6 days (Artific_Inf_Fruits_5_6days), fruits naturally infested in fields sampled after 5-6 days in the lab (Nat_Inf_Fruits_5_6days), fruits naturally uninfested in fields sampled after 5-6 days in the lab (Nat_Uninf_Fruits_5_6days). Fruit Type: blackberry (BL), cherry (CH), strawberry (ST) Fruit sampling site: Thurins (THU), Varetz (VAR), Finestret (FIN), Los Masos (LOM), Ille-sur-Têt (IST), Millas (MIL) Fruit variety: Coeur de Pigeon (CDP), Ciflorette (CIF), Cirafine (CIR), Dream (DRE), Hedelfinger (HED), 80% Mara des bois + 20% Cirafine (INF), Locqtay (LOC), Lochness (LOC), Mara des bois (MDB), Starking (STA), Summit (SUM), Sweetheart (SWE), Unknown (UNK). Unknown_21 bacterial ASV belongs to the Enterobacterales order. Unknown_51 fungal ASV has undetermined Phylum and belongs to the Fungi kingdom.

**Figure S7:**
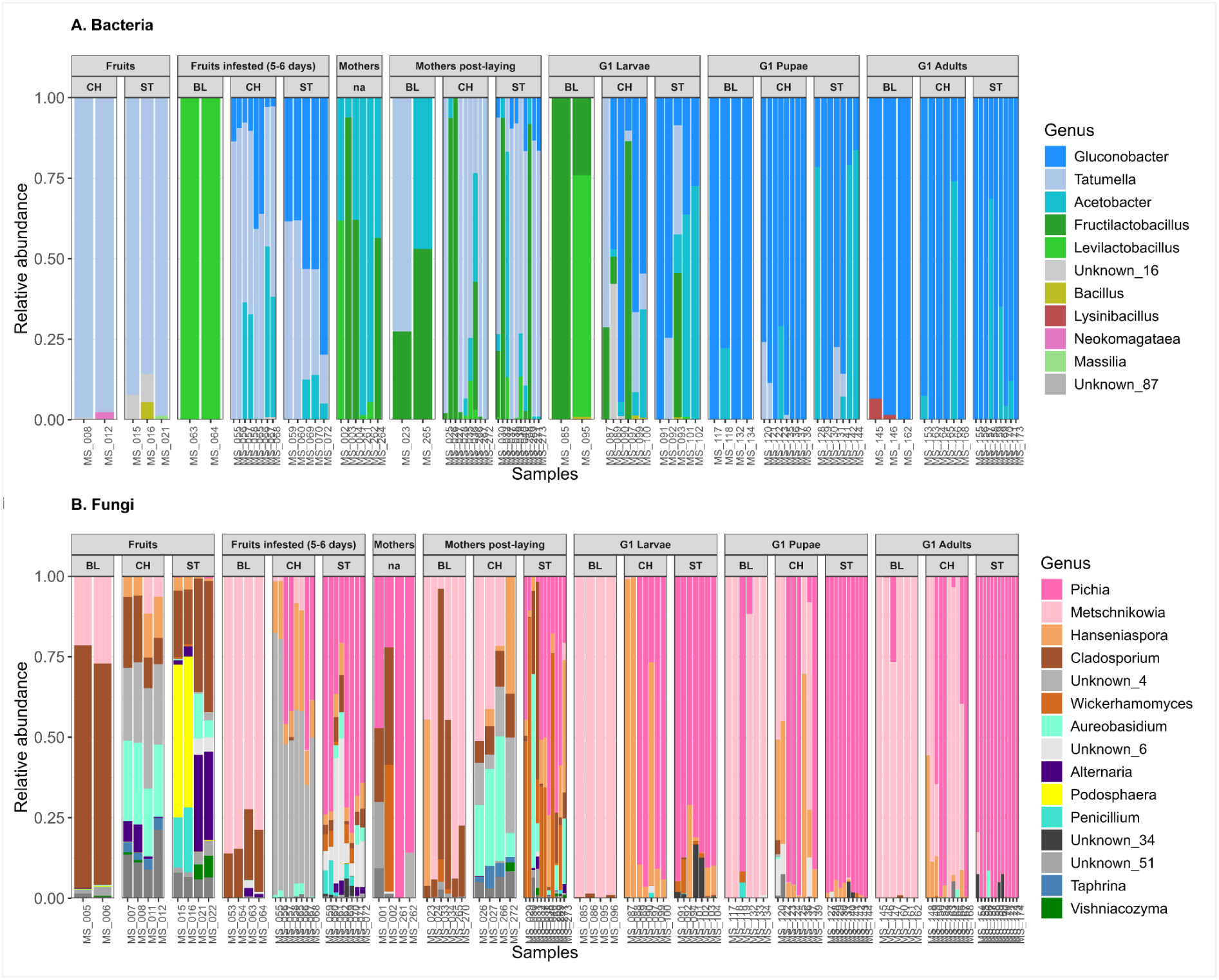
Relative abundance of the 11 most abundant bacterial (A) and 15 fungal genera (B) of *D. suzukii* individuals and their host fruits (Dataset 1), separated by type and host fruit. We sampled undamaged (Fruits) and artificially infested host fruits (Fruits infested 5-6 days), females lab flies used to infest the fruits before (Mothers) and after egg-laying on host fruits (Mothers post-laying), and the offspring at larval (G1 Larvae), pupal (G1 Pupae) and emerged adult (G1 Adults) stages. Host fruit types BL: blackberry, CH: cherry, ST: strawberry Unknown bacterial ASVs at genus level: Unknown_16 belongs to the Erwiniaceae family, and Unknown_87 to Yersiniaceae family. Unknown fungal ASVs at genus level: Unknown_4 and Unknown_6 belong to the Sclerotiniaceae family, Unknown_34 belongs to Saccharomycetales order, and Unknown_51 has undetermined Phylum and belongs to the Fungi kingdom.

**Figure S8:**
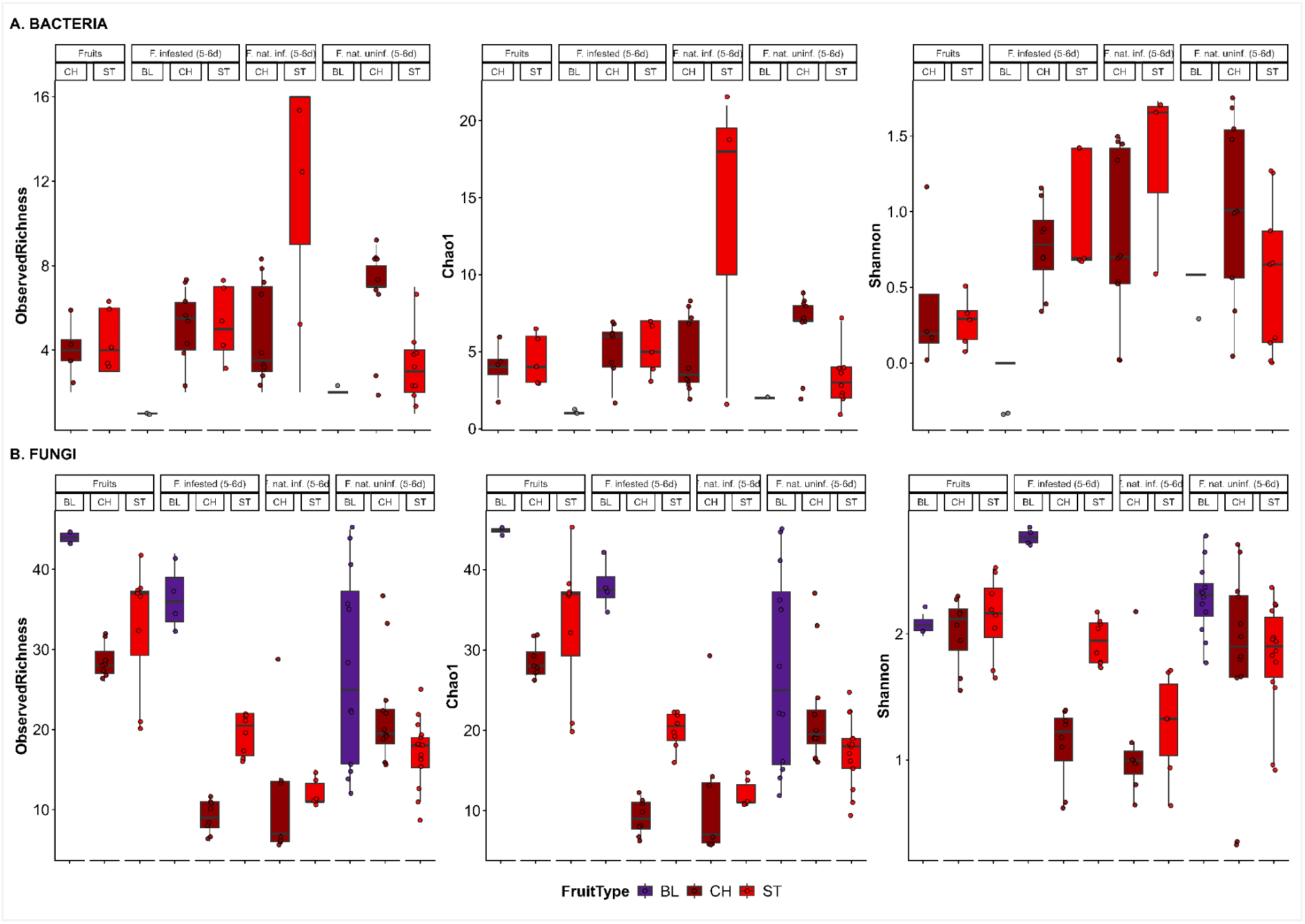
Alpha diversity indices of bacterial (A) and fungal (B) communities present on infested and uninfested fruits, separated by sample type and fruit type (Dataset 2) Sample type: undamaged fruits (“Fruits”), artificially infested fruits (“Fruits infested, 5-6 days”), naturally infested fruits (“Fruits naturally infested, 5-6 days”), and naturally uninfested fruits (“Fruits naturally uninfested, 5-6 days”). Host fruit type: blackberry (BL), cherry (CH), strawberry (ST)

**Figure S9:**
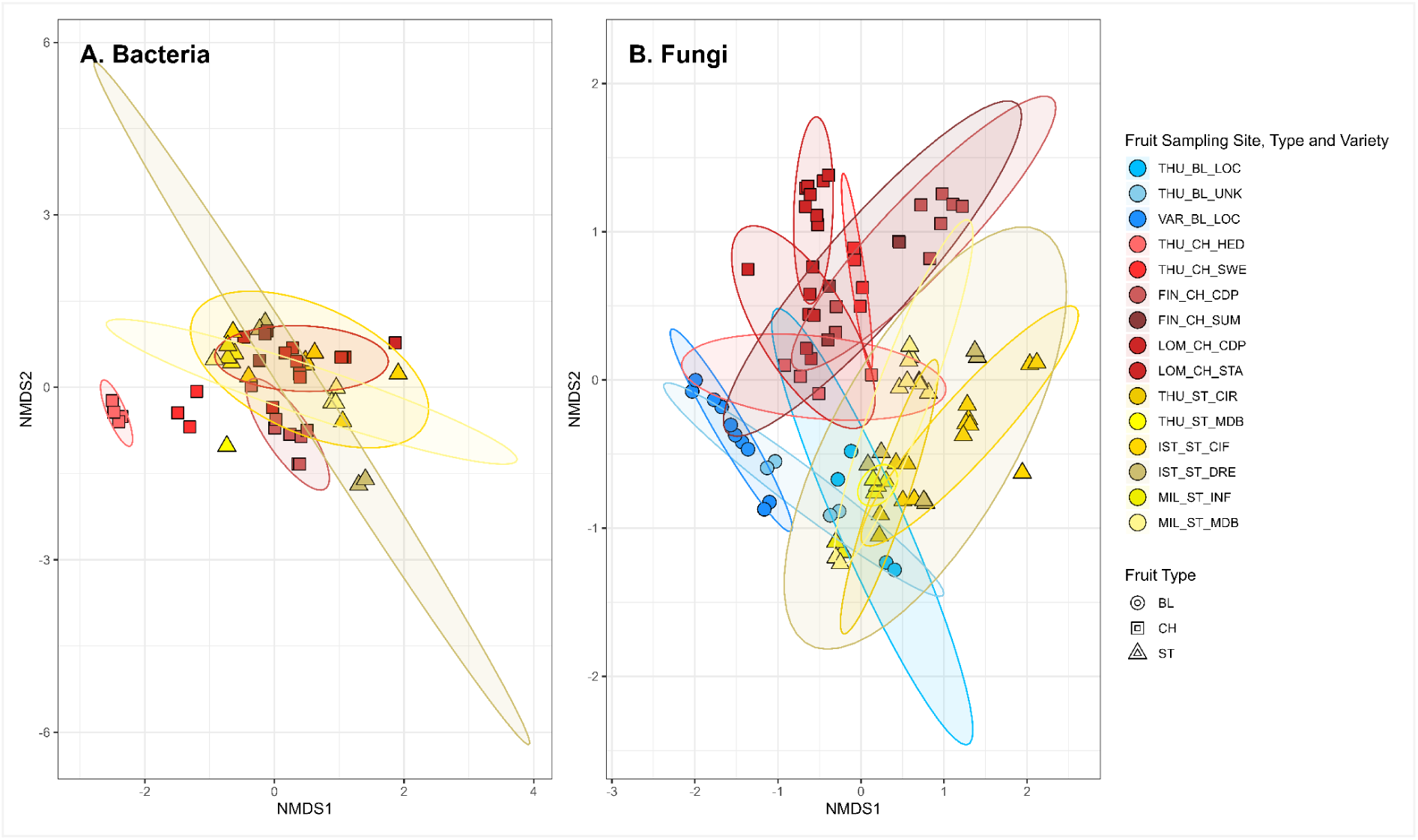
NMDS ordination plots based on Jaccard distances of *D. suzukii* infested and uninfested fruits bacterial (A) and fungal (B) communities (Dataset 2) NMDS ordination plots of bacterial (A) and fungal (B) communities of fruits (stress = 0.1601433 and 0.1894624 respectively). A point represents a pool of fruit (one metabarcoding sample). Symbols represent fruit type: BL = blackberry, CH = cherry, ST = strawberry. Colors represent fruit sampling site x variety: cherries in reddish colors, strawberries in yellowish and blackberries in blueish. Samples ordination based on Bray-Curtis distances was not different, indicating that microbial species’ abundances were balanced (see Fig. S11 in supplemental material).

**Figure S10:**
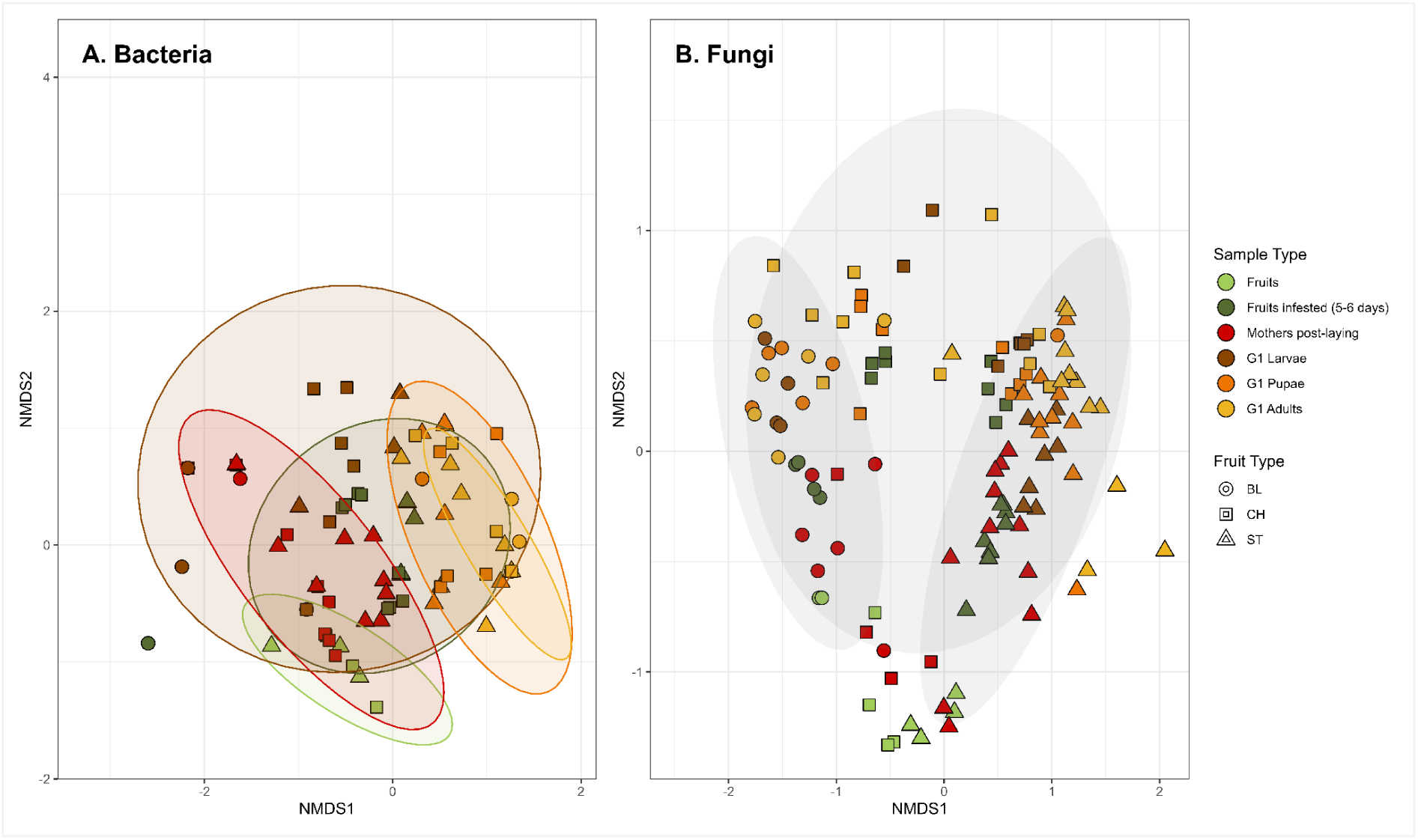
NMDS ordination plots based Bray-Curtis distances of *D. suzukii* individuals and their host fruits bacterial (A) and fungal (B) communities (Dataset 1) NMDS ordination plots based on Bray-Curtis distances of Dataset 1 bacterial (A) and fungal (B) communities (stress = 0.1042379 and 0.1630403 respectively). A point represents a pool of samples (fruits or individuals). Symbols represent fruit type: BL = blackberry, CH = cherry, ST = strawberry. Colors represent sample types. Ellipses are plotted by Sample Type for bacteria (A), and by Fruit Type for fungi (B).

**Figure S11:**
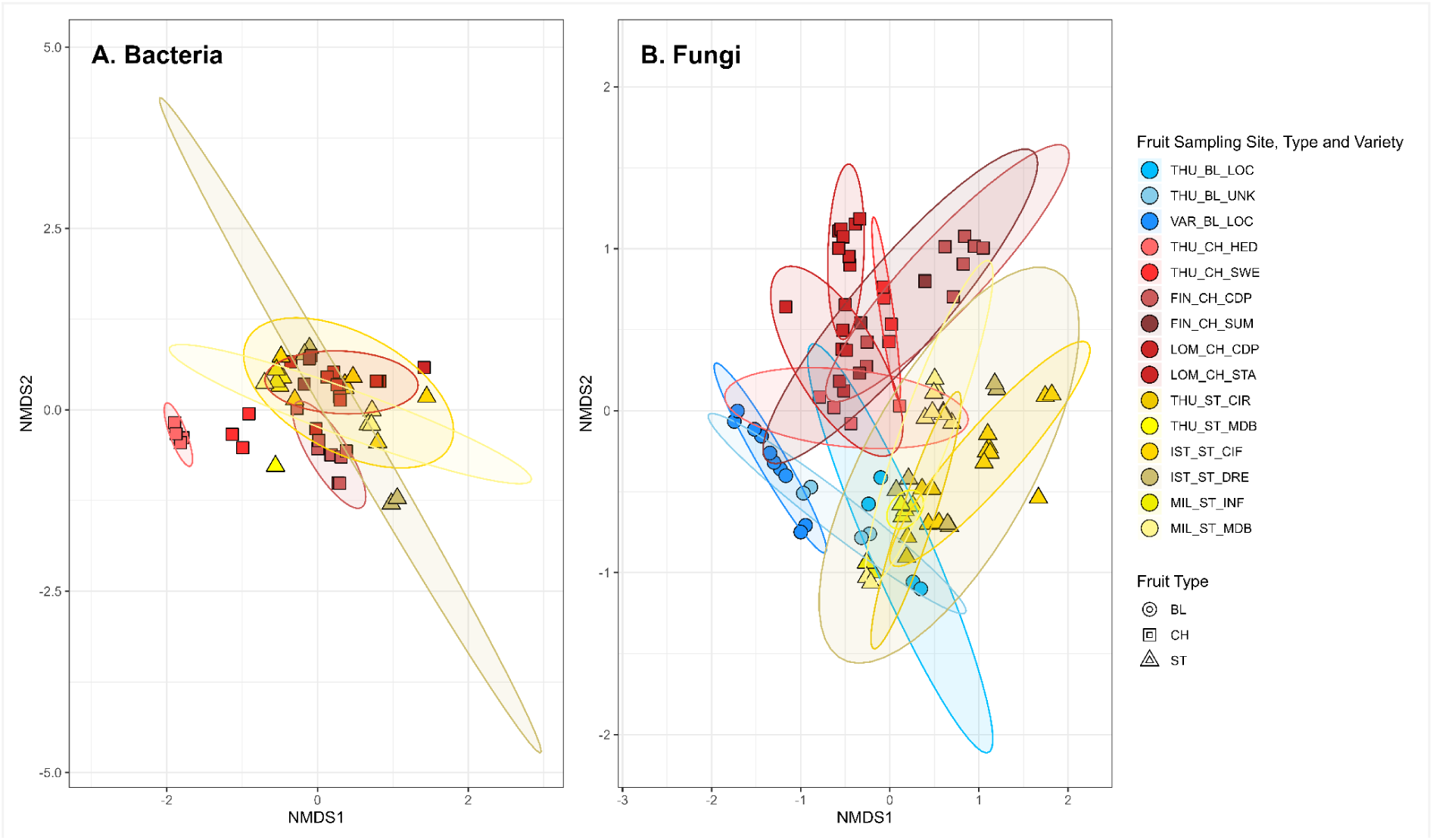
NMDS ordination plots based on Bray-Curtis distances of *D. suzukii* infested and uninfested fruits bacterial (A) and fungal (B) communities (Dataset 2) NMDS ordination plots of host fruit bacterial (A) and fungal (B) communities (stress = 0.1601435 and 0.1894624 respectively). A point represents a pool of fruit (one metabarcoding sample). Symbols represent fruit type: BL = blackberry, CH = cherry, ST = strawberry. Colors represent fruit sampling site x variety: cherries in reddish colors, strawberries in yellowish and blackberries in blueish.

**Figure S12:**
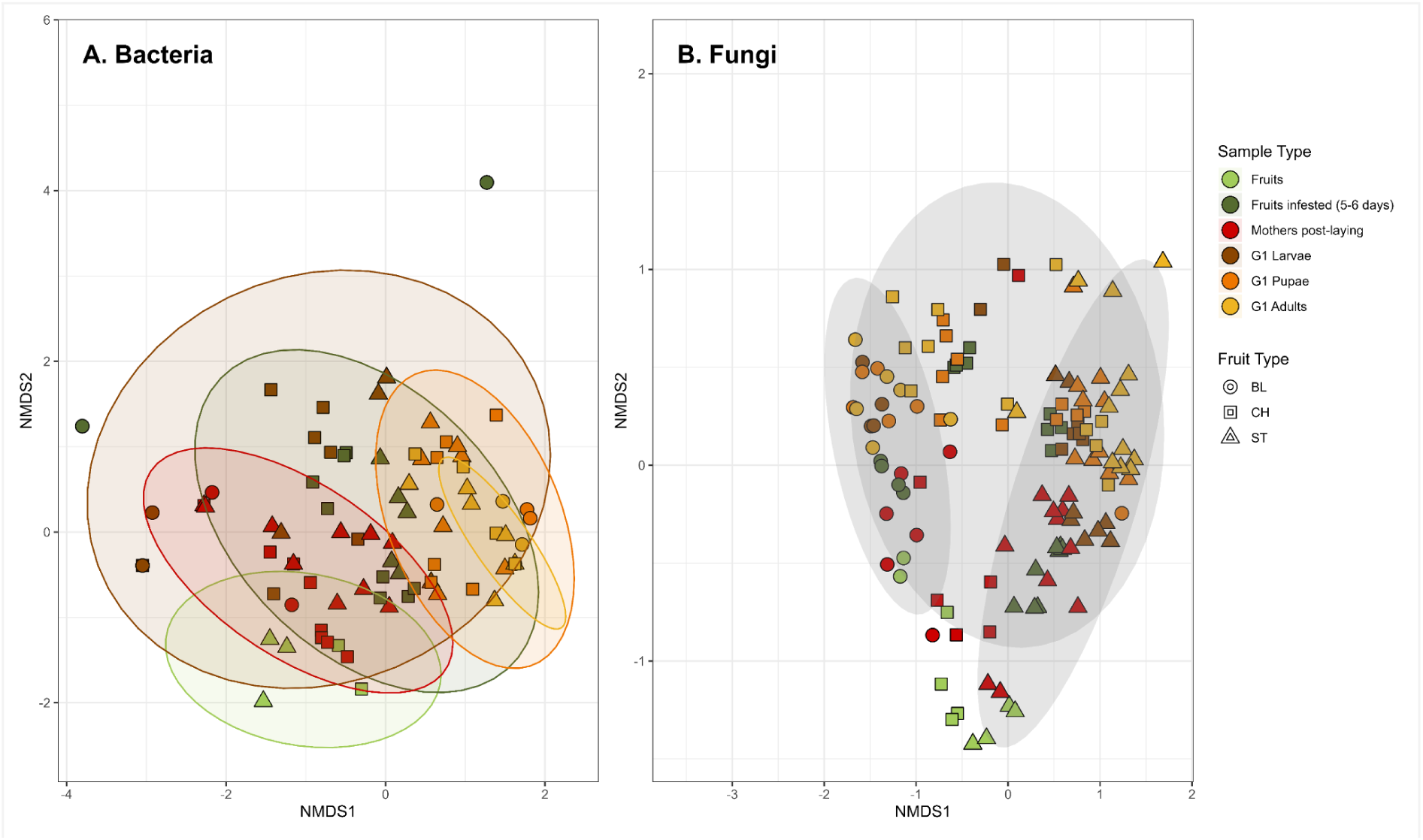
NMDS ordination plots of bacterial (A) and fungal (B) communities of *D. suzukii* individuals and their direct host fruits based on Jaccard distances (Dataset 1), after filtering out ASVs with a prevalence lower than 20%. NMDS ordination plots of *D. suzukii* and its host fruit bacterial (A) and fungal (B) communities (stress = 0.1309201 and 0.175884 respectively). A point represents a pool of samples (fruits or individuals). Symbols represent fruit type: BL = blackberry, CH = cherry, ST = strawberry. Colors represent sample types (mothers before egg-laying samples (Mothers), were pulled out as not related to host fruits). Ellipses are plotted by Sample Type for bacteria (A), and by Fruit Type for fungi (B).

**Figure S13:**
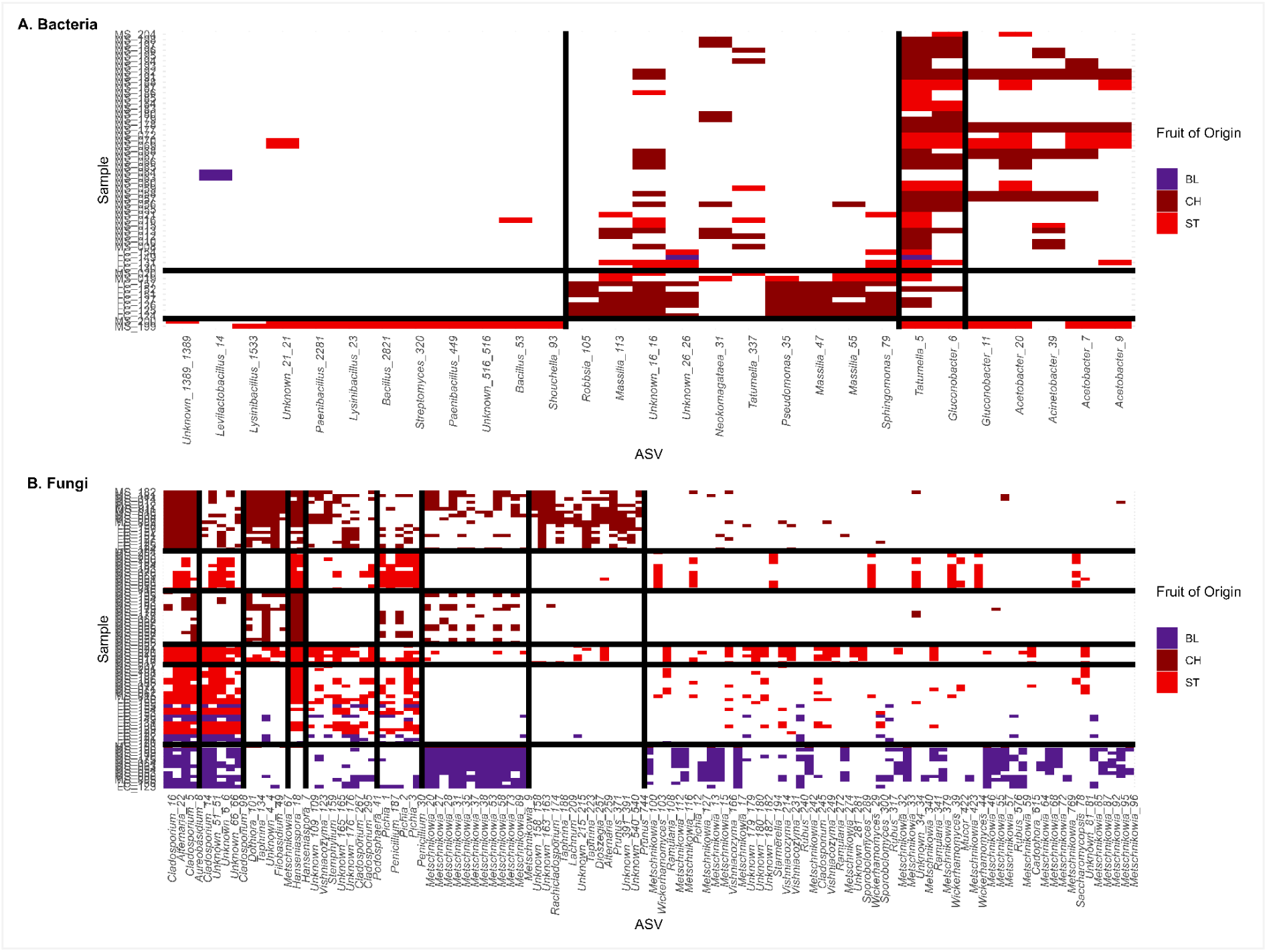
Groups identified from the absence-presence matrix. Infested and uninfested fruits samples were grouped using Latent Block Model (LBM) based on their bacterial (A) or fungal (B) communities (Dataset 2) Presence data colored by fruit type: BL = blackberry, CH = cherry, ST = strawberry. Black lines delimit clusters identified based on the best latent block model.

**Figure S14:**
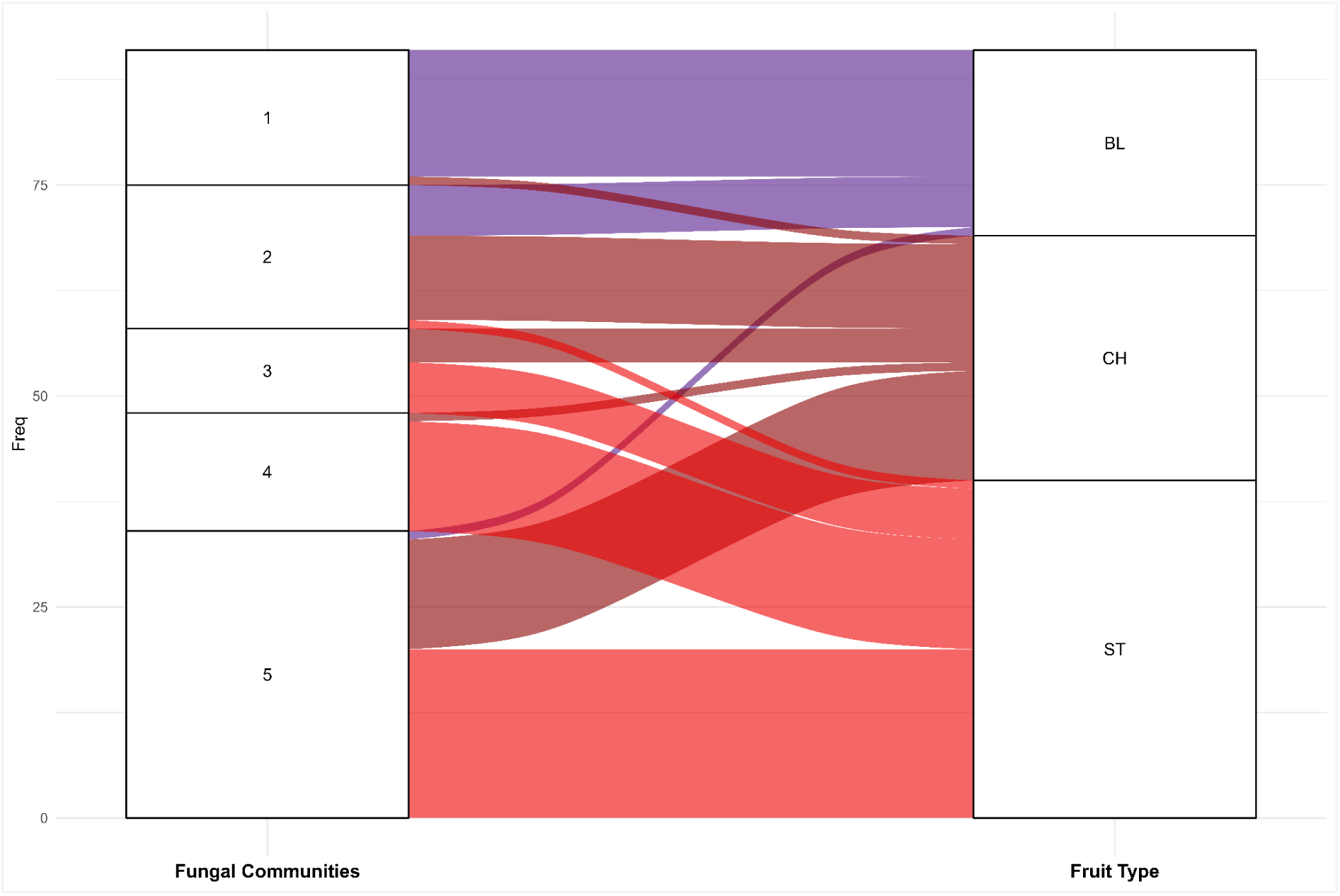
Alluvial plots showing how *D.suzukii* individuals and their host fruits samples group according to fungal communities composition and host fruit type (Dataset 1). Colors correspond to the host fruit type: BL = blackberry, CH = cherry, ST = strawberry; The first grouping factor (Fungal Communities) is the clustering inferred using a latent block model (LBM), based on fungal communities (Fig. 5B), from Dataset 1 presence-absence matrix.

### 2. Supplementary Tables

**Table S1:**
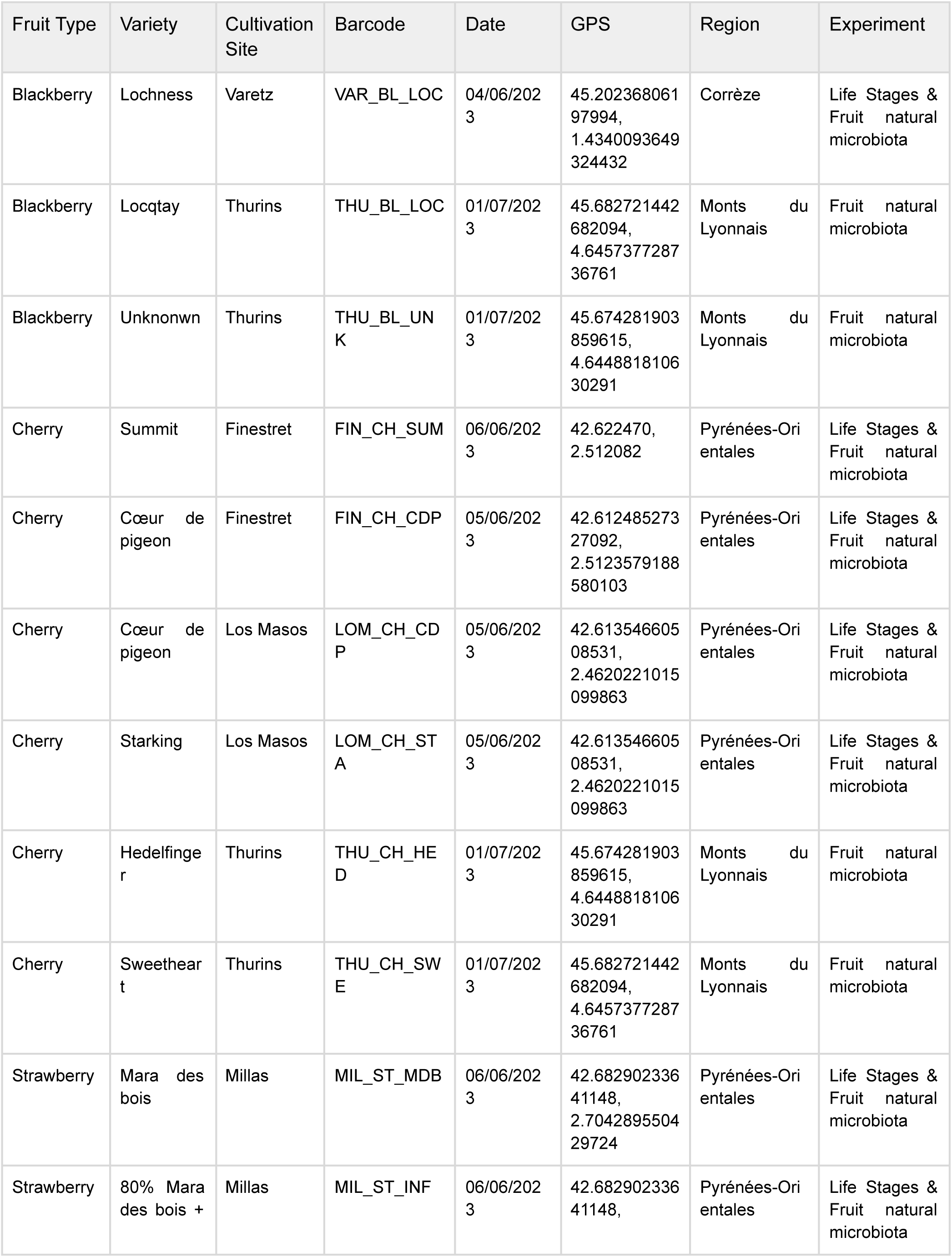
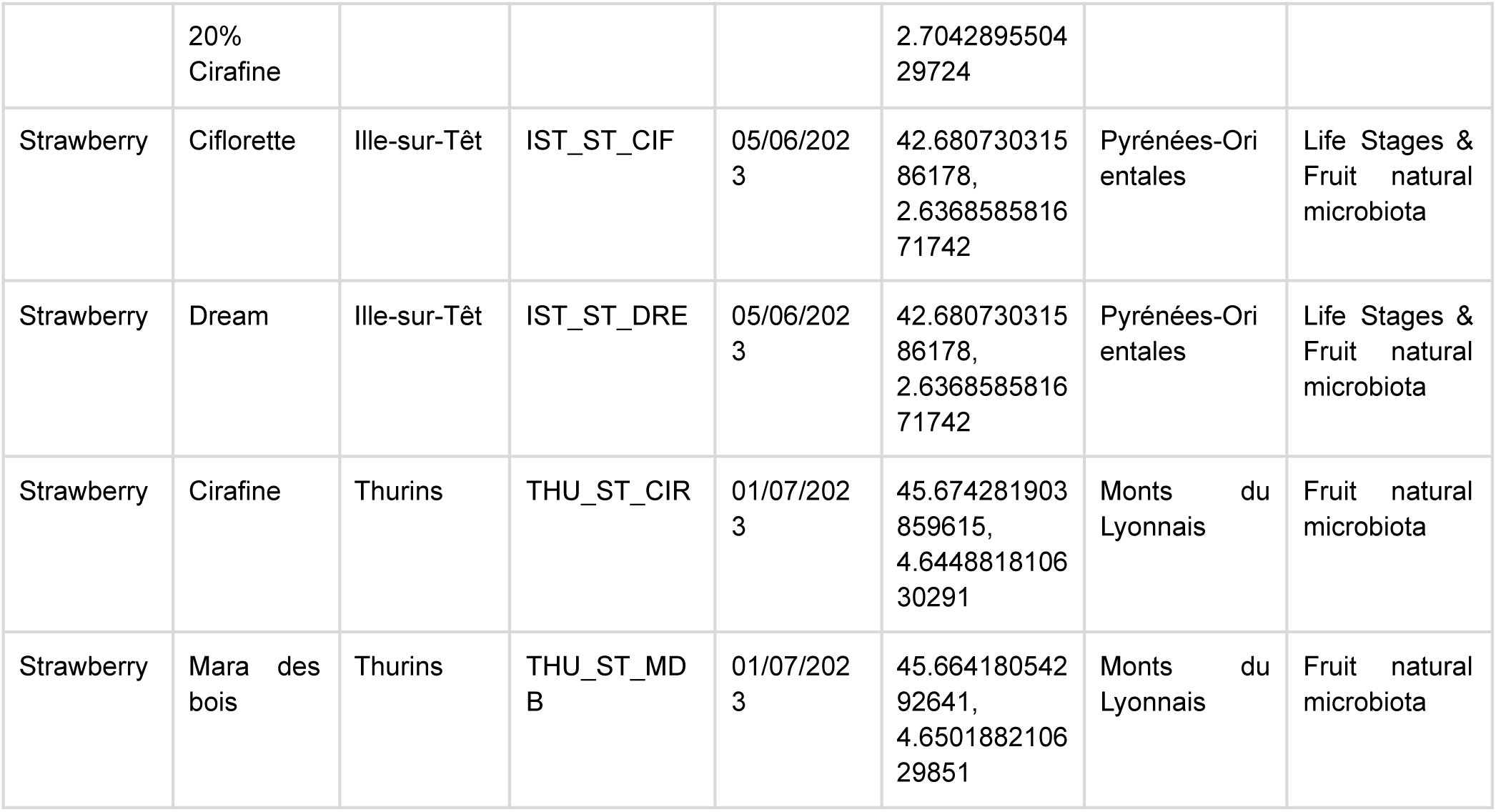
Fruit sampling details.

**Table S2:** reference data for reads proportion and chosen reads repartition.

|  | Fruit_Blackberry | Fruit_Strawberry | Larval | Pupa | Pupal case | Adults |
| --- | --- | --- | --- | --- | --- | --- |
| Bacteria (%) | 13.7 | 5.2 | 37.7 | 90.7 | 95.9 | 99.6 |
| Chloroplasts (%) | 51.8 | 61.7 | 41.8 | 7.3 | 3.5 | 0.0 |
| Mitochondria (%) | 34.6 | 33.1 | 20.5 | 2.0 | 0.0 | 0.0 |

**Reads repartition by markers and sample types chosen for our sequencing**
|  | By_marker_ITS | By_marker_16S | Fruit_16S | Larvae_16S | PupaePupalCases<br>Adults_16S |
| --- | --- | --- | --- | --- | --- |
| Number of reads | 3000000 | 9000000 | 7169541 | 951839 | 878620 |
| Number of samples | 286 | 286 | 68 | 65 | 150 |
| Mean depth | 10489 | 31468 | 5271 | 5418 | 5271 |

**Table S3:** Taxa considered as contaminants and manually removed.

| Contaminant | Taxonomic level considered | Number of clusters removed | Marker |
| --- | --- | --- | --- |
| *Lentihominibacter* | Genus | 1 | 16S |
| *Cutibacterium* | Genus | 14 | 16S |
| *Corynebacterium* | Genus | 86 | 16S |
| Anaerovoracaceae | Family | 1 | 16S |
| Malasseziaceae | Family | 30 | ITS |

**Table S4:** Congruences of between classification of infested and uninfested fruit samples based on bacterial or fungal presence-absence data and fruit type or sample type or sampling site.

| Marker | Factor | NMI | p-value |
| --- | --- | --- | --- |
| 16S | Fruit Type | 0.092 | 0.098 |
| 16S | Sample Type | 0.263 | 0.076 |
| 16S | Fruit Site | 0.346 | 0.1 |
| ITS | Fruit Type | 0.624 | 0.068 |
| ITS | Sample Type | 0.282 | 0.17 |
| ITS | Fruit Site | 0.563 | 0.077 |

Normalized Mutual Information (NMI) Index based on 1000 random permutations
Fruit Type: blackberry, cherry, strawberry. Sample type: undamaged fruits ("Fruits"), artificially infested fruits ("Fruits infested, 5-6 days"), naturally infested fruits ("Fruits naturally infested, 5-6 days"), and naturally uninfested fruits ("Fruits naturally uninfested, 5-6 days").
Fruit Site: Thurins (THU), Varetz (VAR), Finestret (FIN), Los Masos (LOM), Ille-sur-Têt (IST), Millas (MIL)

**Table S5:** CCA between the classification of infested and uninfested fruit samples based on bacterial and fungal presence-absence data and fruit type and sampling site (Dataset 2).

| Formulas | Bacteria |  |  |  | Fungi |  |  |  |
| --- | --- | --- | --- | --- | --- | --- | --- | --- |
|  | chi² | F | p-value (row perm.) | p-value (edge perm.) | chi² | F | p-value (row perm.) | p-value (edge perm.) |
| Fruit Type | 0.1435 | 1.332 | 0.2626 | - | 1.547 | 18.82 | 0.0002 | 0.016 |
| Sampling Site | 0.9720 | 5.080 | 0.001 | - | 2.178 | 12.96 | 0.0002 | 0.0092 |

**Table S6:**
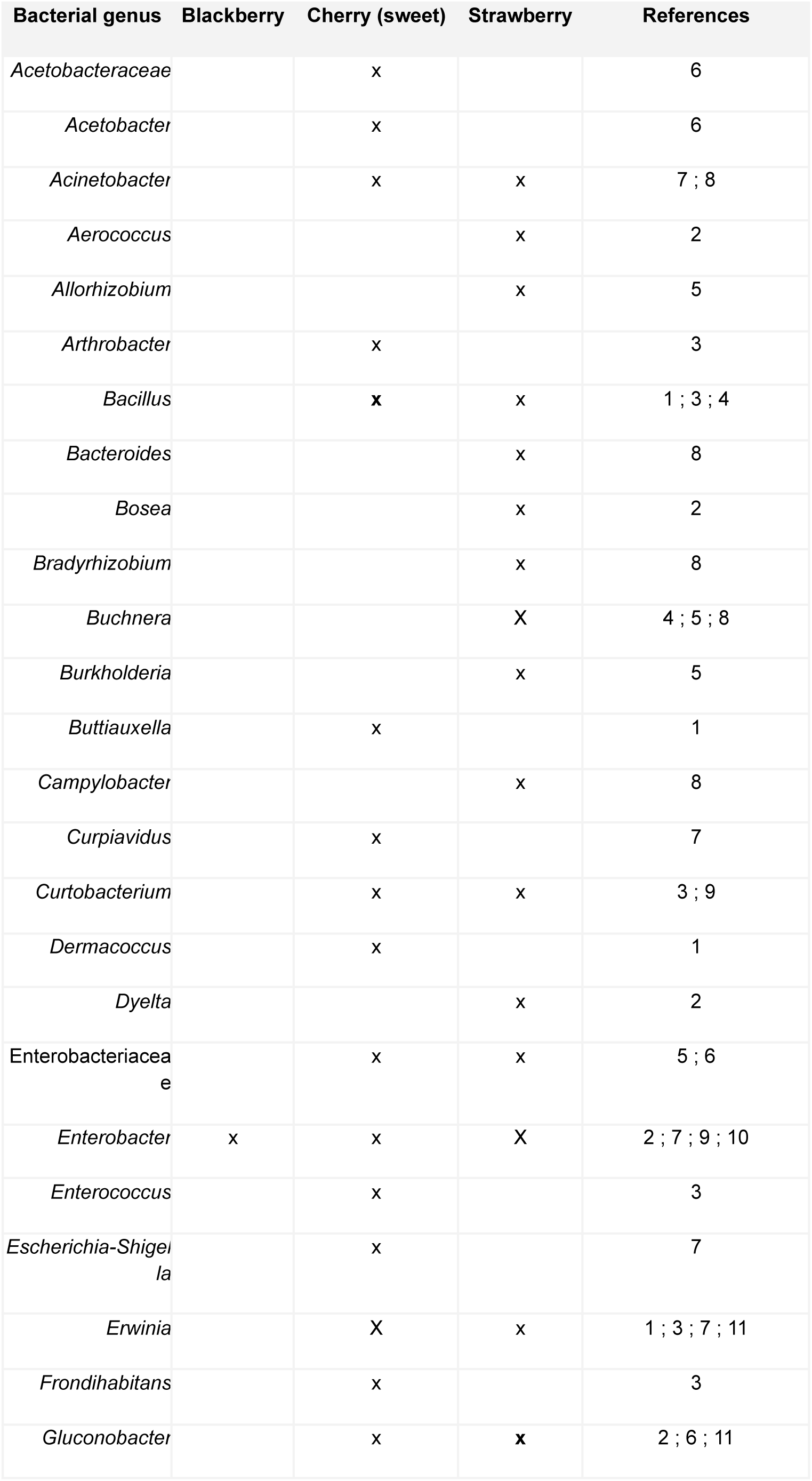
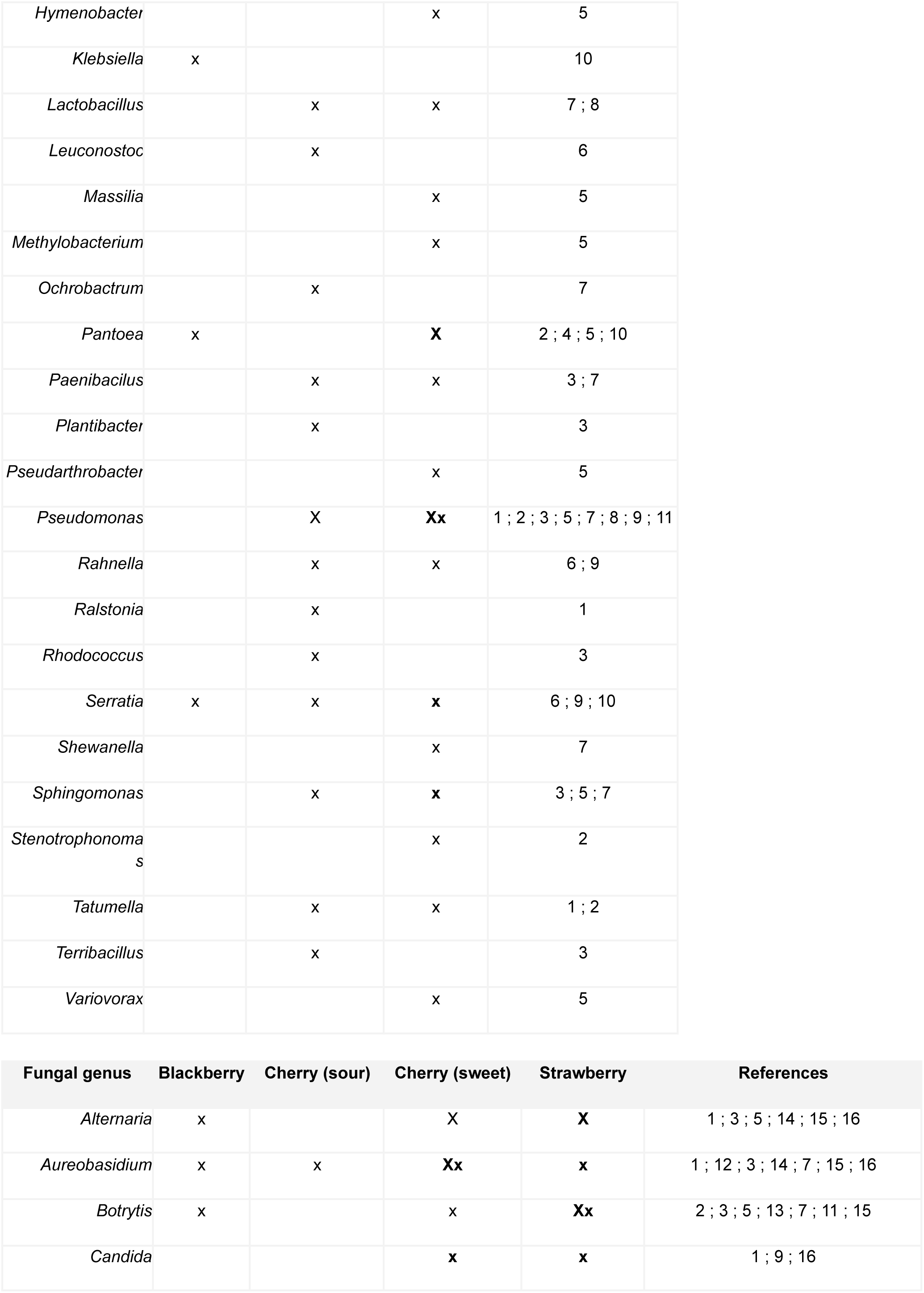
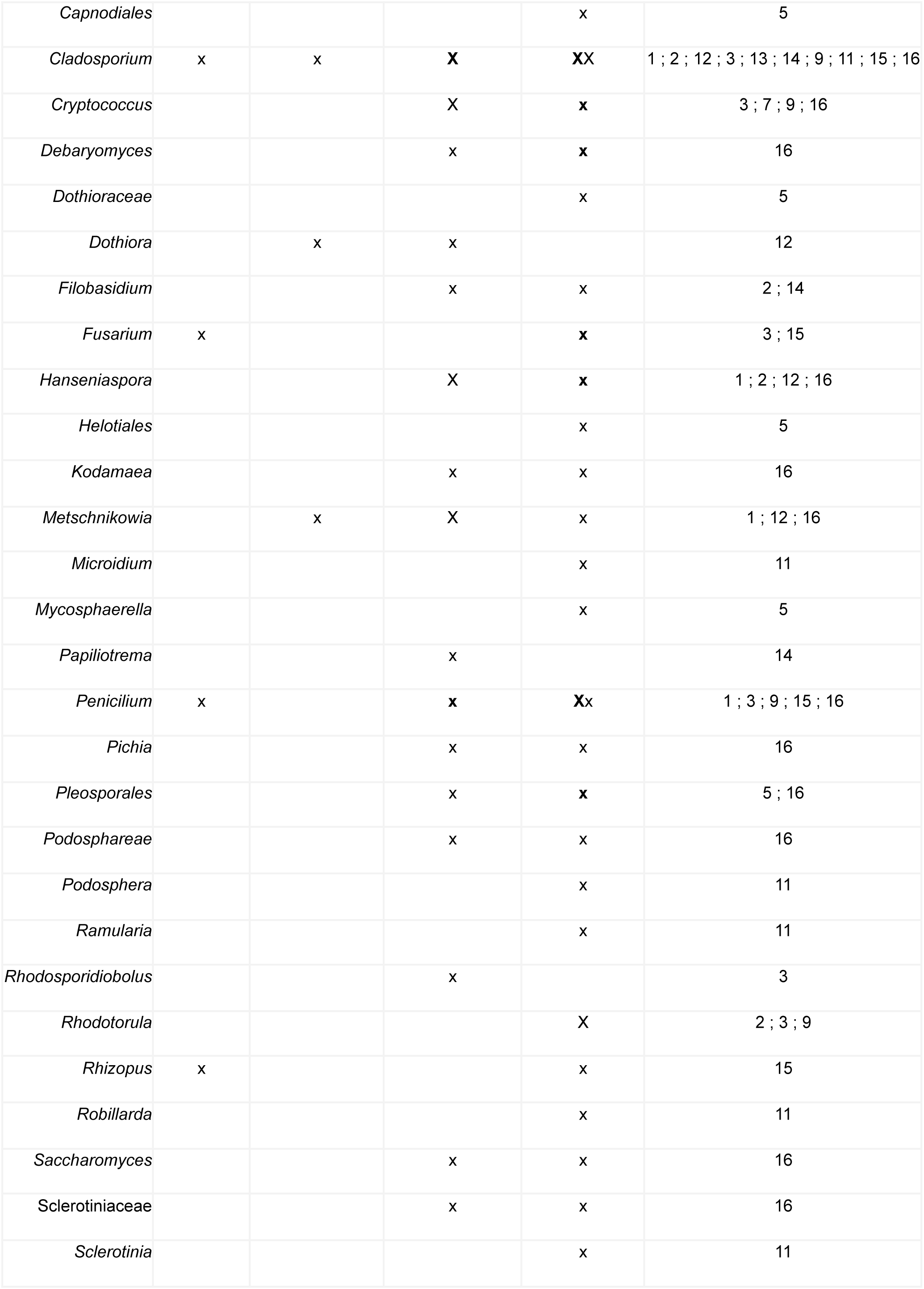
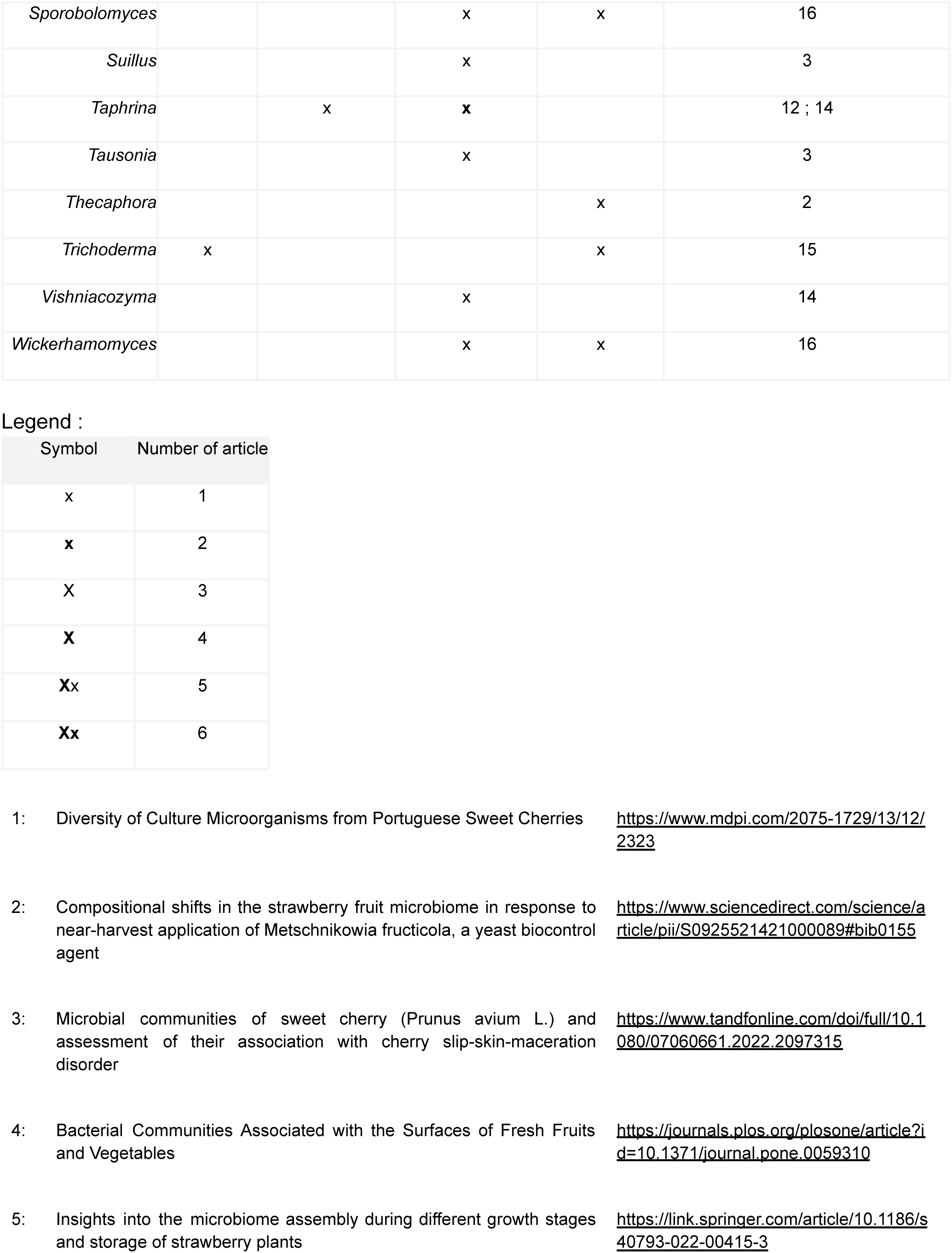
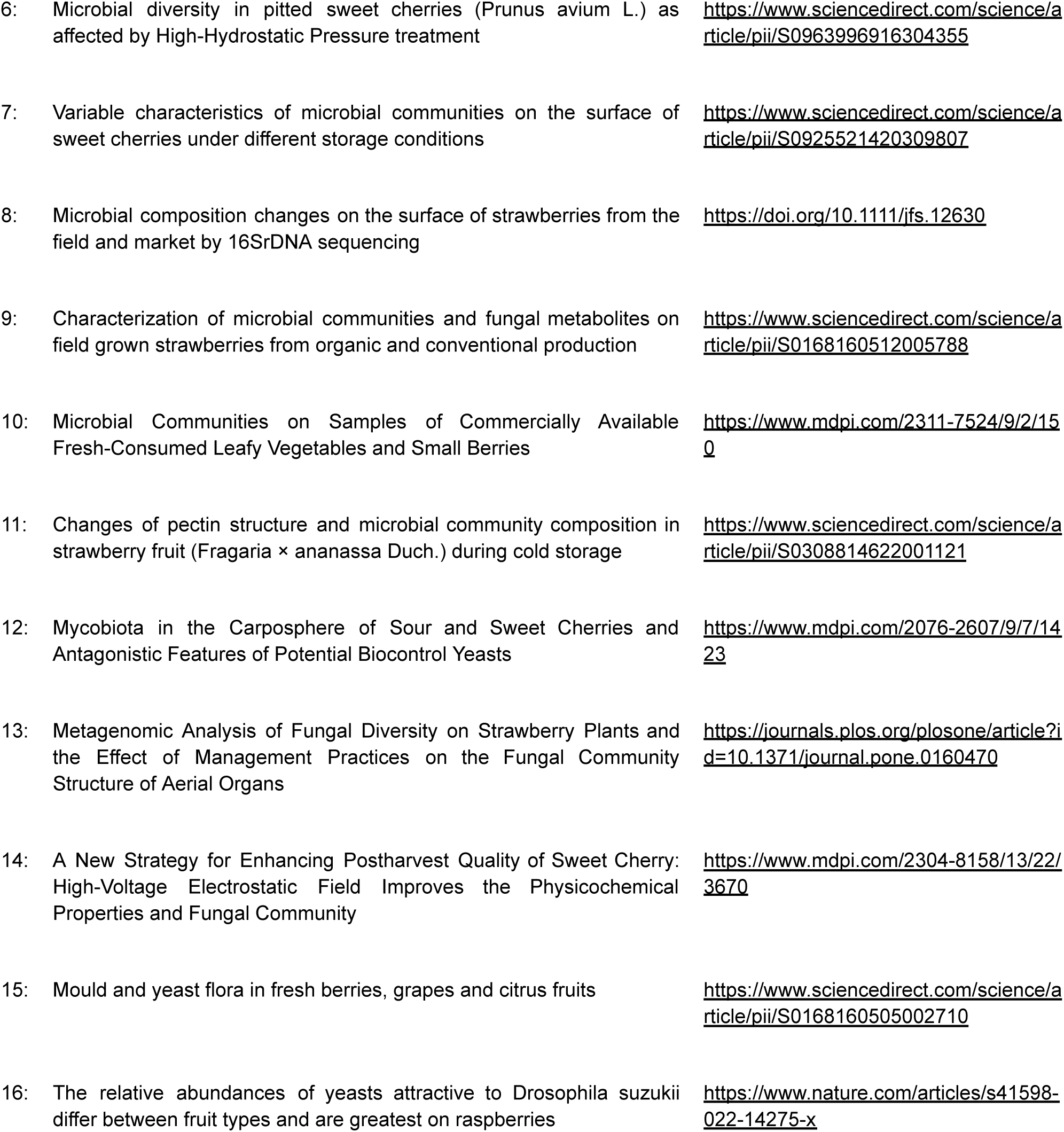
Non-exhaustive review of fruit surface bacterial and fungal species (blackberry, cherry and strawberry).

**Table S7:**
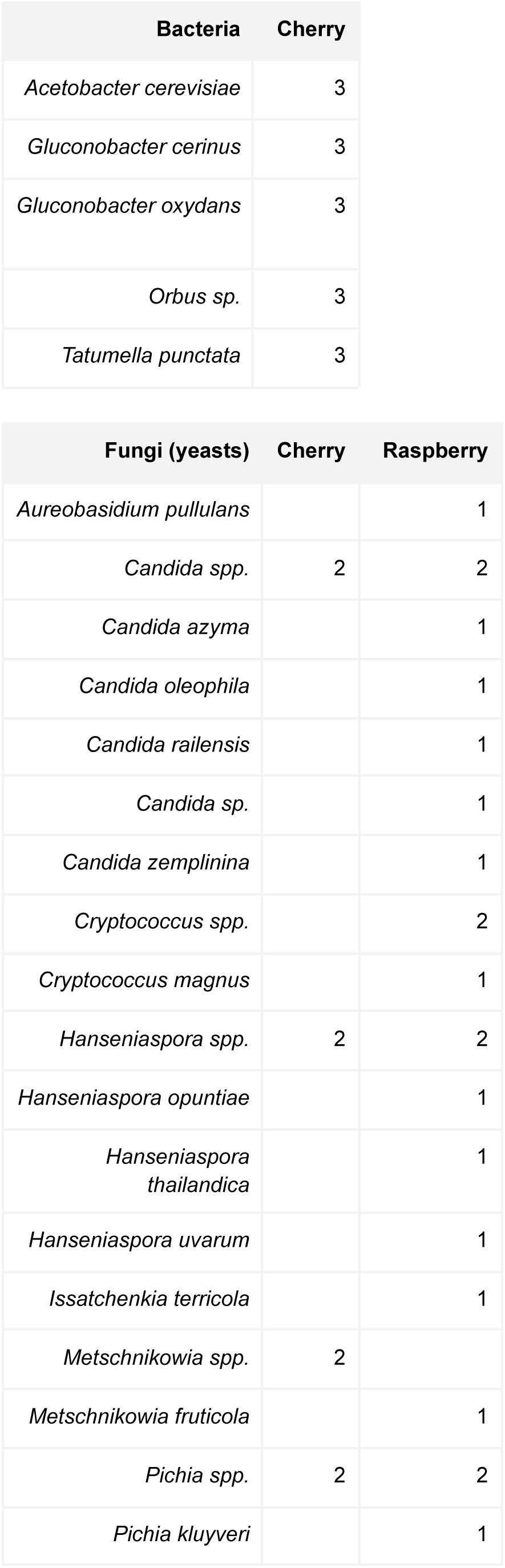
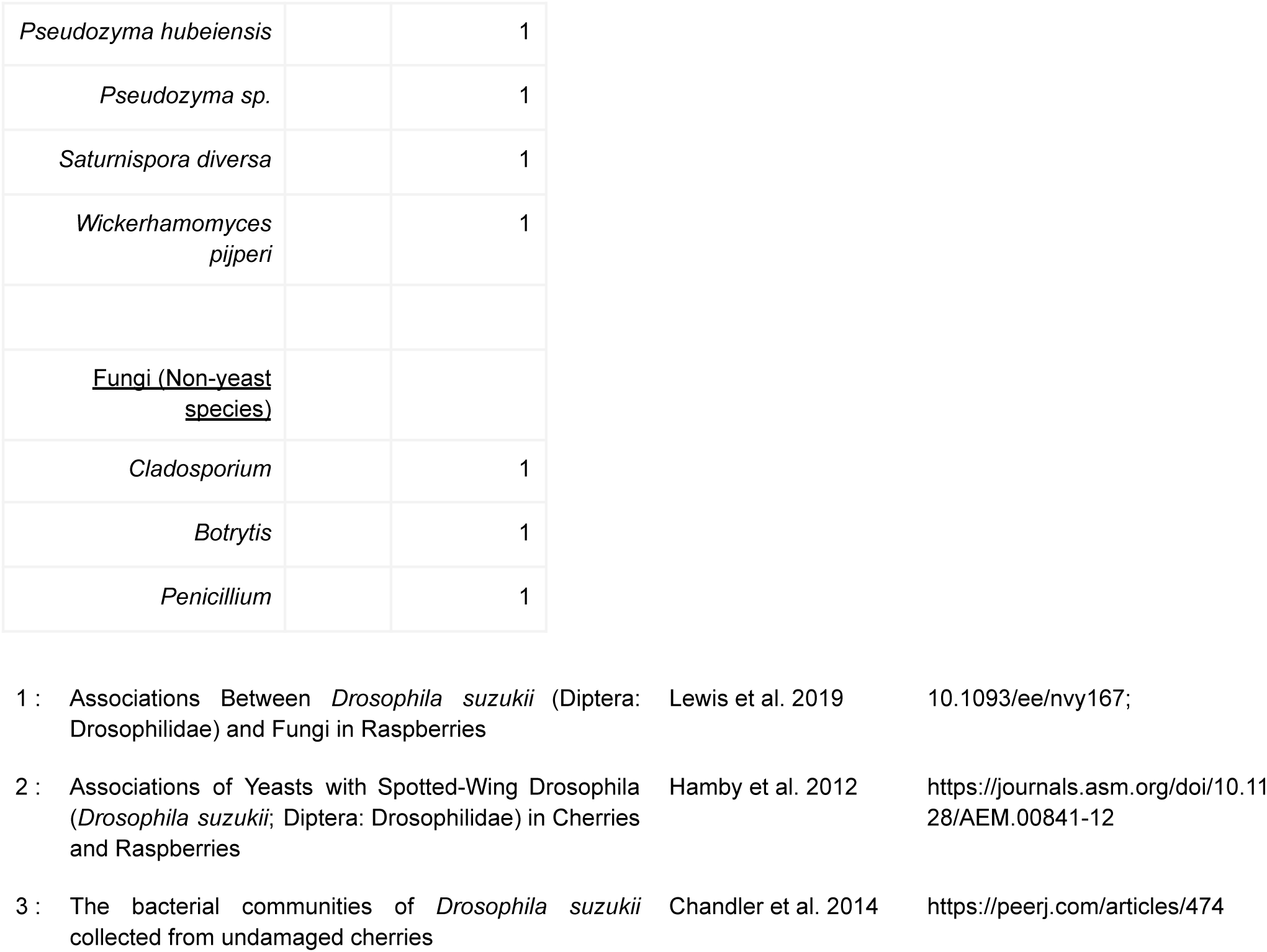
Non-exhaustive review of *D. suzukii* larvae microbiota across host fruits (most frequent species/genera).

### 3. Supplementary Documents

<u>Doc. 1: Controls in the experiment testing for the respective effects of host fruit type and fly life stage on D. suzukii microbiota composition</u>

To compare the compositions of the microbiota of individuals across life stages and between host fruits types, we sampled host fruits, adults, larvae, pupae, pupal cases and emerging adults (Figure 2), and we included different controls.

#### Control of the effect on *D. suzukii* genotype on microbiota composition

To control the effect of the genotype on the flies’ microbiota composition, we used two laboratory populations, bred and maintained separately on cranberry or strawberry medium (CEC or FRA population). At each experimental step, half of the flies were CEC flies and the other half FRA.

#### Fruit microbiota acquisition control

To control if the host fruits were actually the main source of microbes for egg-laying mothers and their offspring, we additionally put mated females on fruit mediums to lay eggs for 48 hours. Fruit mediums were made with pasteurized blackberry, cherry and strawberry purees.

#### Microbiota of larvae from naturally infested fruits

To control host fruit natural infestation for each sampling site and fruit variety, we displayed five infested-looking fruits alone in embryo collection cages during step 1.2 (“Nat_Inf_Fruits_5_6days”). After 5-6 days, during step 1.4 and 1.5, we counted the number of pupae eventually present on the fruit surface and pooled them by five in a sterile 1.5 mL Eppendorf frozen for later metabarcoding. Fruits were transferred individually in a sterile bag after removing the stem or calyx, pressed. We collected 100 µL of the pressed-fruit juice in a sterile 1.5 mL Eppendorf frozen for later metabarcoding. Larvae eventually present in pressed fruits were counted, one-on-two was sampled like pupae (“G1_Larvae_Nat_Inf_Fruits_5_6days”). The other half were individually transferred in a 96-well plate with fruit medium to develop and sampled at pupal and adult stages (“G1_Pupae_Nat_Inf_Fruits_5_6days”, “G1_Adults_Nat_Inf_Fruits_5_6days”) for later metabarcoding. Pupae sampled from fruit surfaces were grouped with pupae developed from larvae on fruit medium (“G1_Pupae_Nat_Inf_Fruits_5_6days”).

#### Naturally non-infested fruit control

In addition, to control the effect of time and of infestation on natural fruit microbiota composition evolution, during step 1.2, we displayed each of ten non-infested-looking fruits in embryo collection cages, for each variety (“Nat_Uninf_Fruits_5_6days”). After 5-6 days, during step 1.5 we controlled the absence of infestation before extracting fruit juice for later metabarcoding as described above. If needed be, we sampled and froze larvae or pupae present and notified natural infestation for the corresponding fruit.

<u>Doc. 2: DNA extraction and PCR amplification detailed protocol</u>

Samples were ground using 3-mm glass beads in ATL lysis buffer solution (DNeasy® 96 Blood & Tissue kit; Qiagen) two times for 2 min at 30 Hz in a tissue lyzer. Total DNA was extracted from tissue homogenates incubated with 20 µL Proteinase K and 180 µL ATL buffer per pool of sample (*i.e.* per microtube) at 55 °C overnight. DNA extraction was continued by grinding samples a second time using zirconium oxide beads (0.45-0.55 mm Zirmil®) two times for 1 min at 30 Hz in a tissue lyzer before following the manufacturer’s instructions (Quick-Start Protocol, DNeasy® 96 Blood & Tissue kit, Qiagen, October 2022) for DNA fixation, wash and elution – except for the final addition of AE elution buffer, for which half of the required volume was used.

The PCR reaction was conducted using 2 µL of sample DNA in 10 µL total volume. The 16S rRNA was amplified using custom-made indexed primers for 16S V4 (16S-V4F 5’-GTGCCAGCMGCCGCGGTAA-3’, 16S-V4R 5’-GGACTACHVGGGTWTCTAATCC-3’), and the ITS rRNA using BITS and B58S3 for ITS1 gene region (BITS 5’-ACCTGCGGARGGATCA-3’, B58S3, 5’-GAGATCCRTTGYTRAAAGTT-3’). Negative PCR and index controls were included to detect PCR problems and index contaminations respectfully. For 16S rRNA amplification, a mock positive control was also included, using Microbial Community DNA Standard (ZymoBIOMICS D6306). The PCR conditions were 15min initial denaturation at 95 °C followed by 35 cycles of denaturation (20 s at 94 °C), annealing (15 s at 55 °C) and elongation (5 min at 72 °C), with a final extension at 72 °C for 10 min.

The PCR amplification of ITS rRNA was performed in two steps. Negative PCR, index controls and a mock positive control were also included, using Mycobiome Genomic DNA Mix (MSA-1010 ™, ATCC®). The PCR1 reaction was conducted using 2 µL of sample DNA in 10 µL total volume. The followed PCR1 protocol was: 15min initial denaturation at 95 °C followed by 37 cycles of denaturation (30 s at 94 °C), annealing (1 min at 52 °C) and elongation (2 min at 72 °C), with a final extension at 72 °C for 10 min. Then indexes and P5-P7 Illumina adaptors were amplified in a second PCR, using UCP Multiplex PCR Master Mix (Qiagen) and 2 µL of sample PCR1 product in 11 µL total volume. The followed PCR2 protocol was: 15min initial denaturation at 95 °C followed by 8 cycles of denaturation (40 s at 95 °C), annealing (45 s at 55 °C) and elongation (2 min at 72 °C), with a final extension at 72°C for 10 min. To check for the presence of fragments of the expected size, the absence of contamination and the homogeneity of concentration between samples, part of PCR products on each plate including all negative and positive controls were run on a 1.5% agarose gel.

PCR products were pooled together by sample type for each marker. The chosen proportions and final volumes taken per sample type within libraries were calculated according to expected sequencing depth based on previous metabarcoding run (see Table S2). Pooled PCR products were run on a 1.25 % agarose gel using « Low Range Ultra Agarose » (BioRad), and bands of the correct size were excised (one for 16S, two for ITS). PCR products in gel bands were purified on columns with NucleoSpin® Gel and PCR Clean-up kit (Machery-Nagel). Excision products were run on a 1.5 % agarose gel to check for residue presence and non-specific bands before dosing the libraries by qPCR (kit Kapa Biosciences). Both 16S and ITS libraries were normalized at 4 nM and pooled in the final library in different proportions. We chose to sur-represent bacterial sequences to compensate for reads taken by mitochondria and chloroplasts (fruit samples), with a ITS /16S ratio at 1 for 4 (see Table S2).

